# Atg23 Interacts With Both the N- and C-termini of Atg9 Via a Hydrophobic Binding Pocket

**DOI:** 10.64898/2025.12.17.694986

**Authors:** Zhanibek Bekkhozhin, Kelsie A Leary, Michael J Ragusa

## Abstract

Macroautophagy is a cellular process where cytosolic material is captured in double membrane vesicles, termed autophagosomes, which fuse with the vacuole or lysosomes leading to the degradation of the captured contents. In yeast, the biogenesis of autophagosomes is initiated by the fusion of a few small vesicles which contain the integral membrane protein Atg9. Atg9 vesicle trafficking is in part regulated by the peripheral membrane protein Atg23. However, the structure of Atg23 and the mechanism by which Atg23 interacts with Atg9 are currently unknown. Therefore, we determined the crystal structure for a monomeric form of Atg23 and characterized the interaction between Atg23 and Atg9. This work reveals that Atg23 contains a novel fold which is consistent with the AlphaFold 3 prediction except that the helices running towards the dimerization region have a bend giving a more curved global architecture than the prediction. In addition, we demonstrate that conserved sequences in both the N and C-terminal regions of Atg9 bind to a hydrophobic cavity on Atg23.

## INTRODUCTION

Autophagy is a set of cellular processes in which cytosolic material is targeted to the vacuole, in yeast, or lysosomes, in higher eukaryotes, resulting in the degradation of this material (1). Autophagy can lead to the degradation of a wide range of cytosolic components including aggregated proteins, organelles and pathogens (2). As such, autophagy is critical for the development, regulation of cellular homeostasis, responding to nutrient starvation, preventing the accumulation of aggregated proteins and also plays important roles in the immune system (3). Due to its role in so many cellular activities, dysfunctions in autophagy have been observed in human diseases including cancer, neurodegenerative diseases, immune disorders and others (4,5). In one example, mutations in the autophagy protein Beclin 1 are observed in 40-75% of sporadic breast, ovarian and prostate cancers and these mutations dramatically increase the rate of lesion formation in mouse models (6).

In yeast, the targeting of cytosolic material to the vacuole can occur via three independent autophagy pathways, termed microautophagy, macroautophagy and chaperone-mediated autophagy (7,8). Macroautophagy, hereafter autophagy, is distinct from the other two autophagic processes in that cytosolic material is captured in large double-membrane vesicles, termed autophagosomes (9). Autophagosomes are generated de novo in the cell initially via the fusion of small 30-50 nm vesicles to generate a short double-membrane sheet termed the phagophore (10–12). The phagophore expands into a cup shape around the cargo to be degraded via lipid transport from the endoplasmic reticulum and potentially via the fusion of additional vesicles (13–17). Once the double membrane is fully expanded around the cargo, it seals leading to the closure of the double membrane and the completion of the autophagosome (18). The outer membrane of the completed autophagosome then fuses with the vacuolar membrane resulting in the degradation of the inner autophagosomal membrane along with the captured contents (19,20). Amino acids and other metabolites are then exported from the vacuole back to the cytoplasm to be reused (21,22).

The biogenesis of autophagosomes requires a set of proteins known as the core autophagy machinery (23). Of the core autophagy proteins, Atg9 is the only integral membrane protein (24,25). Atg9 resides in the small 30-50 nm vesicles that serve as the initial membrane source for phagophore generation (10,11). Atg9 containing vesicles are generated from the Golgi and stored as clusters within the cytosol (11,26). The clusters of Atg9 vesicles then move to autophagy initiation sites which requires the interaction between Atg9 and one of two autophagy initiation scaffolding proteins, Atg17 and Atg11 (27–31). Atg9 also functions as a lipid scramblase to aid in autophagosomal membrane expansion demonstrating that it is both critical for the generation of the phagophore as well as its expansion into the autophagosomal membrane (32,33).

The formation and trafficking of Atg9 vesicles depends on many general vesicular trafficking components, such as soluble N-ethylmaleimide-sensitive (NSF) attachment Receptors (SNAREs), guanosinetriphosphatases, tethering complexes and many others (34–49). In addition to these general vesicle trafficking factors, two more specialized autophagy proteins, Atg23 and Atg27, have been shown to regulate Atg9 vesicle trafficking (11,30,50–54). Atg23 and Atg27 co-localize, co-fractionate and co-precipitate with Atg9 suggesting that these proteins may directly interact with Atg9 to regulate the trafficking of these vesicles (54). Consistent with this, Atg23 deletion blocks exit of Atg9 from Golgi while Atg27 deletion leads to increased Atg9 degradation in the vacuole (10,11,54). As such, Atg23 and Atg27 have been defined as autophagy specific factors that regulate Atg9 vesicle generation and trafficking. However, the exact function and mechanisms of Atg23 and Atg27 are still unknown.

We previously demonstrated that Atg23 is a highly extended alpha helical dimeric protein that can tether vesicles through direct interactions with membranes (55). We also demonstrated that Atg23 dimerization deficient mutants led to reduced Atg9 puncta and reduced recruitment of Atg9 to autophagy initiation sites in yeast further supporting a role for Atg23 in the regulation of Atg9 vesicles biogenesis and/or trafficking. However, a high-resolution structure of Atg23 has not been determined and the molecular mechanism by which Atg23 interacts with Atg9 is still unknown. Therefore, we sought to determine a high-resolution structure of Atg23 and to use this structure along with AlphaFold 3 modeling and biochemical assays to characterize the interaction between Atg23 and Atg9.

## RESULTS

### The generation of monomeric Atg23_ANNS_

We began by attempting to crystalize and determine the structure of dimeric Atg23. However, after extensive crystallization trials with different Atg23 constructs from different yeast species we were unable to obtain crystals that diffracted with sufficient resolution for structure determination. Instead, we investigated the AlphaFold 3 predicted structure of dimeric Atg23 to determine if smaller fragments of Atg23 might be more amenable for experimental structure determination (56). The predicted structure of Atg23 is predominantly helical with a maximum dimension close to 300 Å, which is consistent with our previous small angle X-ray scattering envelope and modelling (Figure 1A) (55). Atg23 is predicted to contain nine α-helices and a single β-strand (Figure 1B and S1). All α-helices, except for α6 are predicted with pLDDT confidence scores > 70 indicating that these elements are predicted with confidence (Figure 1C). In contrast, pLDDT scores for α6 are all < 70 indicating that this helix is predicted with low confidence. Atg23 is also predicted to contain a C-terminal disordered tail which spans approximately amino acids 382-453.

**Figure 1.**
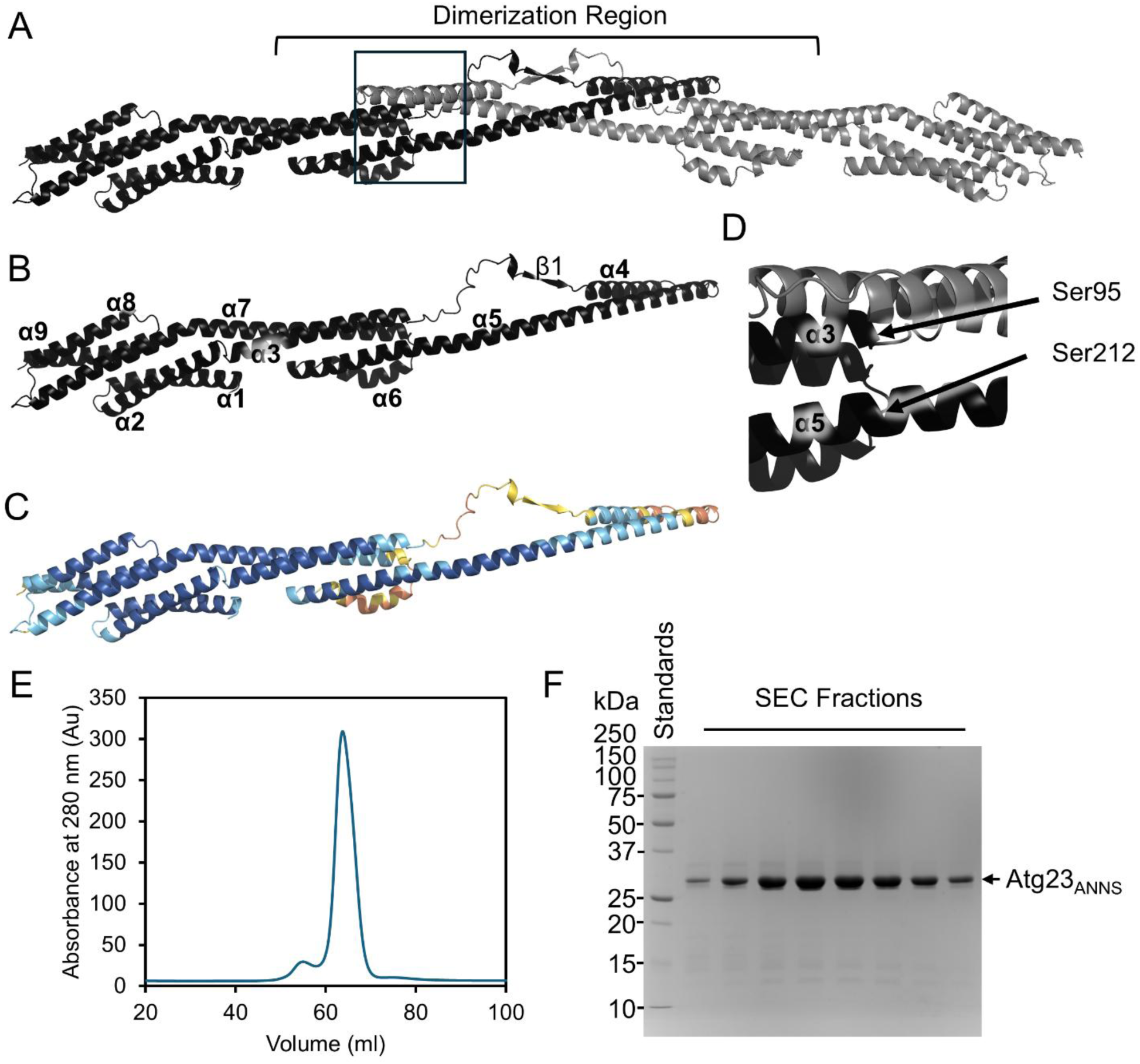
AlphaFold structure prediction guided design of Atg23_ANNS_. **A)** Cartoon representation of the AlphaFold 3 predicted structure of the Atg23 dimer. One monomer is shown in black and the other in grey. The flexible disordered region is now shown for clarity. **B)** One monomer of the Atg23 dimer from A is shown as a cartoon representation with each element of secondary structure labeled. **C)** One monomer of Atg23 is shown with the confidence pLDDT scores colored dark blue for very high (plDDT > 90), blue for confident (90 > plDDT > 70), yellow for low (70 > plDDT > 50) and orange for very low (plDDT < 50). **D)** A close-up view of the Atg23 dimer (boxed in A) highlighting the proximity of Ser95 and Ser212. **E)** SEC chromatogram of Atg23_ANNS_. **F)** SDS-PAGE gel showing the purity of Atg23_ANNS_ as the protein eluted from the column in D.

Dimerization of Atg23 appears to be supported by an extended three helical bundle comprising both copies of α5 from each monomer and α4 from a single monomer (Figure 1A). A single beta sheet containing one beta-strand from each monomer also appears to play a role in mediating dimerization although the pLDDT scores for this region are not as confident as they are for α4 and α5 (Figure 1C). Intriguingly, this dimerization region is formed by the middle region of Atg23, spanning amino acids 95 to 212. This suggests that dimerization is encoded in the middle of the protein and that dimerization cannot be removed via simple truncation at either terminus of Atg23 while leaving the remainder of the protein intact for structural analysis. α5 is the largest helix in Atg23 and it partially participates in dimerization but also has intramolecular interactions with α3 of the same monomer. Further investigation of the structure revealed that the dimerization interface appears to end after Ser212 in α5. Ser212 is in close proximity to Ser95 from α3 (Figure 1D). Therefore, we reasoned that it might be possible to remove the entire dimerization region of Atg23 and create a short linker between these two amino acids to generate a stable monomeric form of Atg23. This construct would lack amino acids 96 to 211, including all of α4 and β1 and most of α5 which is predicted to form the majority of the dimer interface.

To generate a construct of Atg23 that might be amenable to crystallization we replaced amino acids 96 to 211 in Atg23 with a short four amino acid linker (Ala-Asn-Asn-Ser) which we predicted would be long enough to connect Ser95 to Ser212 without imposing any new structural restraints on the protein (Figure 1D). We also removed the C-terminal tail spanning residues 382-453 as this region is predicted to be disordered and would likely prevent crystallization. This construct, which we termed Atg23_ANNS_, expressed well in *E. coli* (∼200 mg protein from 1L of culture) and could be purified to high purity and concentrated to 20 mg/ml. Atg23_ANNS_ eluted from size exclusion chromatography as a single sharp peak suggesting that the protein is monodisperse and soluble (Figure 1E and 1F). Dynamic light scattering further confirmed that the protein was monodisperse (Figure S2A). Circular dichroism of Atg23_ANNS_ demonstrated that the protein was predominantly helical which is consistent with the predicted structure of Atg23 (Figure S2B). Interestingly, Atg23_ANNS_ had a melting temperature of 55°C compared to dimeric Atg23 with 43°C suggesting that Atg23_ANNS_ is more thermostable than dimeric Atg23 (Figure S2C).

### The crystal structure of monomeric Atg23_ANNS_

Atg23_ANNS_ crystalized in 0.07M MOPSO, 0.03M Bis-Tris, 8.8% PEG-8000, 17.6% 1,5-pentanediol and with the monosaccharides II additive which included 20 mM each of xylitol, myo-inositol, D-(-)-fructose, L-rhamnose, D-sorbitol. Atg23_ANNS_ produced long rod-like crystals in the P2_1_ space group that diffracted to 2.1 Å (Table 1). We generated selenomethionine containing Atg23_ANNS_ and collected a single-wavelength anomalous diffraction (SAD) dataset for phasing. However, the anomalous signal was too weak to obtain accurate phases. Instead, we combined SAD phasing and molecular replacement using the most globular distal part of Atg23 including amino acids 1-65 and 288-381 of the full-length Atg23 dimer AlphaFold 3 model. We had attempted to use the complete AlphaFold 3 model of Atg23_ANNS_ for molecular replacement phasing (56). However, this was not successful likely due to structural differences between the predicted and experimental structures. The majority of the electron density map was clear and enabled us to build most of the protein with the exception of the amino acids between α5 and α7 (Fig S3A and S3B). We also observed very weak helical density near where we would expect α6 to be located based on the AlphaFold 3 prediction (Fig S3C). However, we could not see any side chain density and so we built this helix as a series of unknown (UNK) amino acids to show its location in the structure but to not attribute any specific amino acids to this helix. The poor density in this region compared to the rest of the structure is consistent with the possibility that this region of the protein is more dynamic than the rest of Atg23. The low pLDDT scores for this region, amino acids 239-259, in the AlphaFold 3 model of dimeric Atg23 are also consistent with possible structural heterogeneity and conformational dynamics in this region.

**Table 1.** X-ray crystallography data collection and refinement statistics.

|  |  |
| --- | --- |
| **Space group** | P2<sub>1</sub> |
| **a, b, c [Å]** | 50.41, 30.10, 91.24 |
| **$\alpha$, $\beta$, $\gamma$ [°]** | 90.0, 97.15, 90.0 |
| **Resolution range [Å]** | 46.29-2.10 (2.16-2.10) |
| **Completeness [%]** | 100 (100) |
| **Multiplicity** | 3.8 (3.9) |
| **$\langle I/\sigma(I) \rangle$** | 5.6 (1.5) |
| **R<sub>merge</sub>** | 0.115 (0.966) |
| **R<sub>pim</sub>** | 0.103 (0.831) |
| **CC<sub>1/2</sub>** | 0.990 (0.653) |
| Refinement |  |
| **Unique reflections used** | 16327 (1546) |
| **R<sub>work</sub>/R<sub>free</sub>** | 0.238/0.298 |
| **No. protein atoms** | 2001 |
| **No. waters** | 100 |
| **Mean B values for protein [Å<sup>2</sup>]** | 53.06 |
| **Mean B values for water [Å<sup>2</sup>]** | 61.15 |
| **RMSD for bonds [Å]** | 0.003 |
| **RMSD for angles [°]** | 0.531 |
| **Ramachandran favored/allowed [%]/outliers [%]** | 96.7/2.5/0.8 |

The overall structure of Atg23_ANNS_ is consistent with the AlphaFold 3 predicted structure of Atg23 except that α3 and α5 have a different angle pointing towards where the dimerization region would be in full-length Atg23 (Figure 2A). When the Atg23_ANNS_ structure is aligned to α1, α2, α8 and α9 of the Atg23 AlphaFold 3 model, these regions of the structure align well along with the majority of α7 (Figure 2B). However, α3 and α5 are shifted down by approximately 12 degrees (Figure 2B). When the Atg23_ANNS_ structure is instead aligned to the ends of α3 and α5 in the Atg23 AlphaFold 3 predicted structure there is a significant shift in the position α1, α2, α8 and α9 suggesting that the structure of Atg23, while similar to the Atg23 AlphaFold 3 predicted structure, might have a different overall shape (Figure 2C). We next generated a hybrid structural model of dimeric Atg23 by combining our X-ray crystal structure of Atg23_ANNS_ with the AlphaFold 3 predicted structure. We replaced amino acids 1-95 and 212-382 with the Atg23_ANNS_ structure. This hybrid structural model of Atg23 reveals that Atg23 likely contains a more curved architecture than the AlphaFold 3 predicted structure of the Atg23 dimer (Figure 2D and 2E).

**Figure 2.**
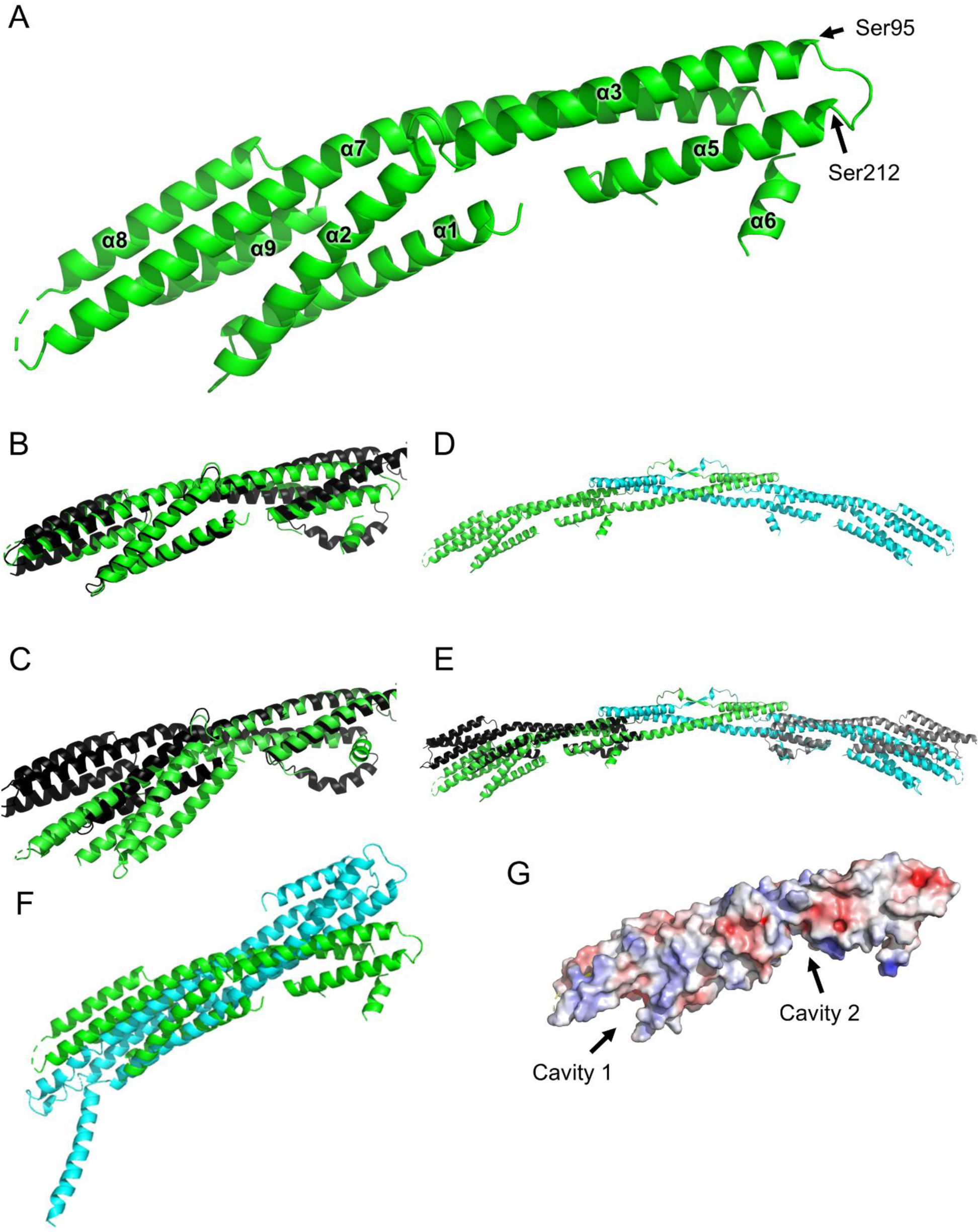
The crystal structure of Atg23_ANNS_. **A)** Cartoon representation of the Atg23_ANNS_ crystal structure. Each element of secondary structure is labeled corresponding to the labeling from the Atg23 dimer. Ser95 and Ser212 are also labeled. **B)** Overlay of the Atg23_ANNS_ (green) with one monomer of the Atg23 dimer (black) aligned based on α1, α2, α8 and α9. **C)** Overlay of the Atg23_ANNS_ (green) with one monomer of the Atg23 dimer (black) aligned based on α3 and α5. **D)** Hybrid structural model of the Atg23dimer with Atg23_ANNS_ replacing the corresponding region if each Atg23 monomer. **E)** Overlay of the Atg23 hybrid structure (green and cyan) from D with the AlphaFold 3 predicted structure of the Atg23 dimer (black and grey). These structures were aligned based on the dimerization region of Atg23. **F)** Comparison of Atg23_ANNS_ (green) with the top hit, cytolysin A pore from *Escherichia coli* (cyan, PDBID: 2WCD), from a Foldseek search. **G)** Electrostatic surface representation of Atg23 where blue is positively charged and red is negatively charged. The two largest cavities which are potential protein binding sites are labeled.

To determine if Atg23 is structurally similar to other proteins we performed a Foldseek search that compares the sequence and tertiary structure of an input model, in this case Atg23_ANNS_, to a defined set of structural databases (57). We restricted our search to the Protein Data Bank (PDB) to compare Atg23_ANNS_ to experimentally determined structures. To our surprise, the top Foldseek hits were coiled-coil containing proteins with no clear connection to membrane binding or vesicle biogenesis. The top two hits were the cytolysin A pore from *Escherichia coli* (PDBID: 2WCD) and a truncation mutant of a dynein motor domain from *Dictyostelium discoideum* (PDBID: 3VKG). The RMSD between Atg23_ANNS_ and these structures is 13.07 Å and 24.42 Å, respectively, indicating poor similarity between these structures. Both structures visibly align poorly to Atg23_ANNS_ further suggesting that the Atg23_ANNS_ fold is not similar to any previously determined structures (Figure 2F). Given that Atg23 may interact directly with Atg9 in addition to membranes we analyzed the surface of Atg23_ANNS_ for potential protein-protein interaction surfaces using the Cavity Plus Server (58). Atg23_ANNS_ contains two primary cavities with volumes of 1161.62 Å^3^ (cavity 1) and 647.38 Å^3^ (cavity 2) which could be potential binding sites for small molecule ligands or other proteins including Atg9 (Figure 2G).

### Atg23 interacts with the N-terminal tail of Atg9

Atg9 forms a trimer in the membrane with each monomer containing four membrane spanning helices and two helices partially embedded in the membrane (24,32,33). Both termini of Atg9 are predicted to be intrinsically disordered, which has been experimentally verified for the N-terminal region of *S. cerevisiae* Atg9 using NMR spectroscopy (Figure 3A) (29). The N-terminal region of Atg9 was also shown to directly interact with Atg11 via two conserved PLF motifs facilitating its trafficking to autophagy initiation sites (29). Given this, we began by asking if dimeric Atg23 can bind to the intrinsically disordered N-terminus of Atg9 spanning residues 1-255. We first used biolayer interferometry (BLI) which demonstrated that Atg9 1-255 binds to Atg23 with a dissociation constant (K_d_) of 1.5 ± 0.7 µM (Figure 3B, S4A and Table 2). Binding was also verified using isothermal titration calorimetry (ITC) which gave a K_d_ of 5.6 ± 1.9 µM and an n of 0.80 ± 0.17 (Figure S4B and Table 3). Interestingly, BLI reported a K_d_ that is 3.7 times tighter than ITC. This could be due to the difference in techniques since in BLI Atg9 is restricted on the surface of a tip potentially adding an avidity effect to this interaction that was not observed in ITC.

**Figure 3.**
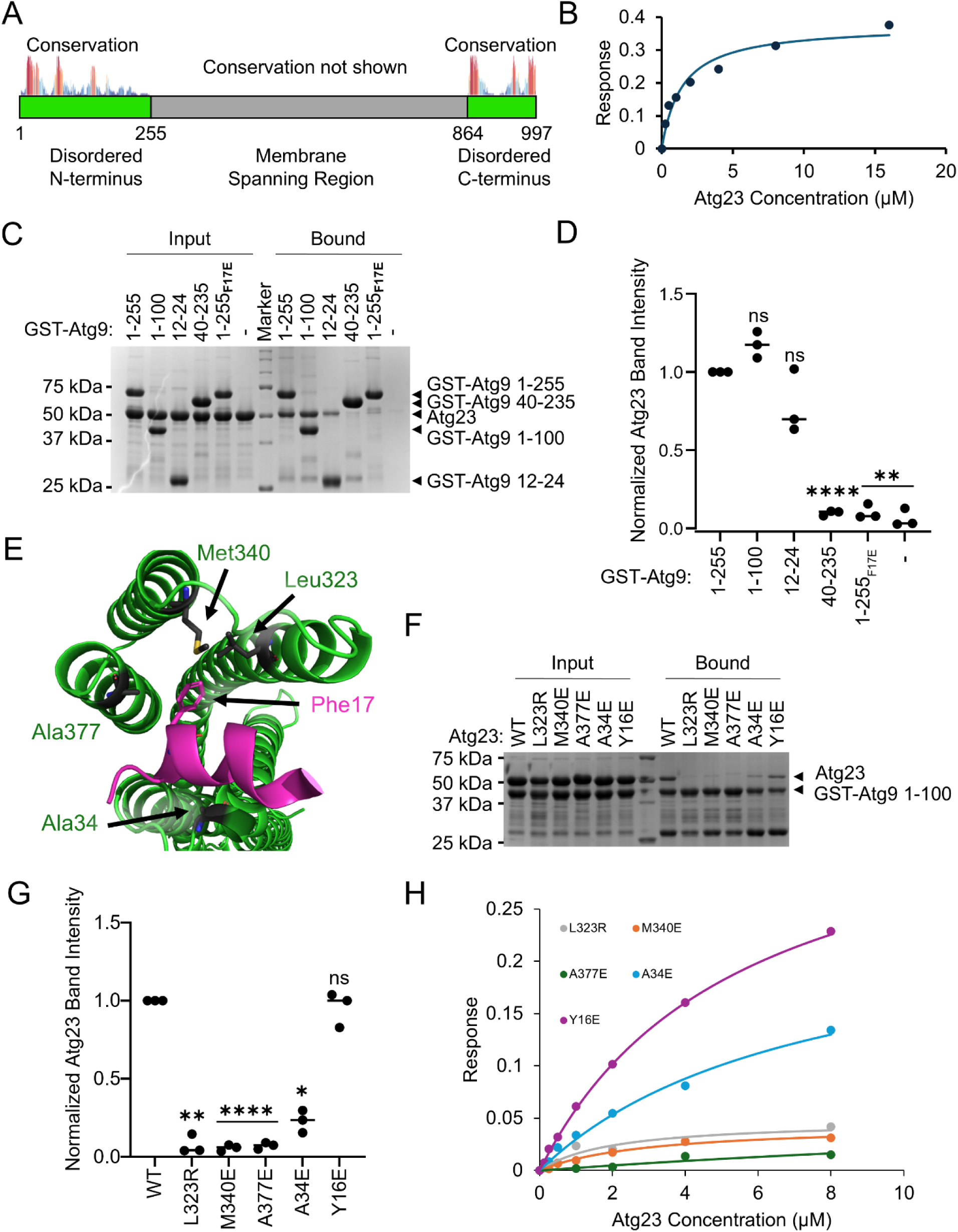
Atg23 interacts with the N-terminal region of Atg9. **A)** Domain architecture of Atg9. The intrinsically disordered regions of Atg9 are colored in green while the membrane spanning region is grey. Amino acid sequence conservation of the intrinsically disordered regions is indicated above the domain architecture where red indicates highly conserved and blue indicates partially conserved. Conservation is not shown for the membrane spanning region. **B)** Steady state analysis of the BLI data for Atg9 1-255 and Atg23. Data is shown as dots and the fit to the data is shown as a line. **C)** SDS-PAGE gel for the GST-Atg9 pulldowns of Atg23. The molecular weight (MW) of each marker is shown on the left and the identity of each band is labeled on the right. **D)** Quantification of three independent repeats of C. Each replicate is shown as a dot and the average is shown as a line. **E)** Cartoon representation of the AlphaFold 3 model of Atg9 1-100 interaction with Atg23. Atg9 amino acids 14-25 (magenta) are shown as these are the amino acids that interact with Atg23. Atg9 Phe7 and Atg23 Ala34, Leu323, Met340 and Ala377 are all shown as stick representations and labeled. **F)** SDS-PAGE gel for the GST pulldown of Atg9 1-100 with different Atg23 point mutants. The MW of each marker is shown on the left and the identity of each band is labeled on the right. **G)** Quantification of three independent repeats of F shown as in D. **H)** Steady state analysis of the BLI data for Atg9 1-255 and different Atg23 point mutants. Data is shown as dots and the fit to the data is shown as a line. Statistical analysis in D and G was performed as a One-way ANOVA with data in D being compared to GST-Atg9 1-255 and data in G being compared to WT. **** P ≤ 0.0001, ** P ≤ 0.01, * P ≤ 0.05, ns, not significant.

**Table 2.** Dissociation constants (K_d_) calculated from BLI data. His6-Atg9 1-255, His10-GST-Atg9 864-997 and His10-GST-Atg9940-997 with Atg23 were repeated in triplicate while the remainder of the samples were performed once to validate GST-pulldowns. n.d. indicates not determined and was reported for samples that had binding close to background levels and therefore a K_d_ could not be determined for these samples.

| Sample on Tip | Sample Added | $K_d$, $\mu\text{M}$ |
| --- | --- | --- |
| His6-Atg9 1-255 | Atg23 | $1.50 \pm 0.7$ |
| His6-Atg9 1-48 | Atg23 | 1.23 |
| His6-Atg9 1-255 | Atg23 L323R | n.d. |
| His6-Atg9 1-255 | Atg23 M340E | n.d. |
| His6-Atg9 1-255 | Atg23 A377E | n.d. |
| His6-Atg9 1-255 | Atg23 A34E | 7.4 |
| His6-Atg9 1-255 | Atg23 Y16E | 5.8 |
| His10-GST-Atg9 864-997 | Atg23 | $0.46 \pm 0.13$ |
| His10-GST-Atg9 940-997 | Atg23 | $0.37 \pm 0.13$ |
| His10-GST-Atg9 940-997 | Atg23 L323R | n.d. |
| His10-GST-Atg9 940-997 | Atg23 M340E | n.d. |
| His10-GST-Atg9 940-997 | Atg23 A377E | n.d. |
| His10-GST-Atg9 940-997 | Atg23 A34E | 0.57 |
| His10-GST-Atg9 940-997 | Atg23 Y16E | 1.72 |

**Table 3.** Thermodynamic data from ITC experiments. Atg9 samples were added to the cell and Atg23 was titrated into Atg9. Each experiment was repeated three times and the values are reported as the mean with standard deviation.

| Cell | Syringe | $K_d$ ($\mu\text{M}$) | N | $\Delta H$, $\text{kJ mol}^{-1}$ | $\Delta S$, $\text{J mol}^{-1} \text{K}^{-1}$ |
| --- | --- | --- | --- | --- | --- |
| His6-Atg9 1-255 | Atg23 | $5.6 \pm 1.9$ | $0.8 \pm 0.17$ | $-7.4 \pm 0.1$ | $75.6 \pm 2.4$ |
| Atg9 864-997 | Atg23 | $0.79 \pm 0.12$ | $0.79 \pm 0.2$ | $-13.7 \pm 1.6$ | $70.4 \pm 5.1$ |

We examined the sequence conservation of the N-terminal region of Atg9 to identify regions of high conservation that might be possible binding regions for Atg23 (Figure 3A). We compared 23 different yeast Atg9 sequences and found that the regions with highest sequence conservation in the N-terminal disordered region are 15-42, 69-84 and 135-146. Guided by the sequence conservation we purified a series of GST tagged Atg9 constructs, 1-255, 1-100, 12-24 and 40-235 to test which region of Atg9 is interacting with Atg23. We performed a pull-down assay where different constructs of GST-Atg9 were bound to glutathione resin, followed by addition of Atg23 and a resin wash. The amount of Atg23 remaining bound to the resin after the wash was quantified. We observed that GST-Atg9 1-100 had similar amounts of Atg23 binding as GST-Atg9 1-255 suggesting that the first 100 amino acids of Atg9 are critical for Atg23 binding (Figure 3C and 3D). Furthermore, GST-Atg9 12-24 had slightly reduced but still significant binding of Atg23 while GST-Atg9 40-235 had almost no Atg23 binding (Figure 3C and 3D). Given that Atg9 40-255 had no binding for Atg23 we purified Atg9 1-48 and checked its interaction with Atg23. We found that Atg9 1-48 bound to Atg23 with a K_d_ of 1.23 µM suggesting that residues 1-48 in Atg9 contain the entire Atg23 binding region (Figure S4D and Table 2).

We next used AlphaFold 3 to predict the molecular interaction between Atg9 1-48 and Atg23 (Figure 3E). Atg9 16-26 formed an alpha helix and bound directly into cavity 1 on Atg23. Atg9 17-19 were predicted with confident pLDDT scores while the remainder of Atg9 was predicted with low or very low pLDDT scores suggesting that amino acids 17-19 might be the most critical for binding. Atg9 Phe17 contributed the largest amount of buried surface from Atg9 and was centered in the hydrophobic binding pocket of cavity 1 surrounded by Atg23 Leu323 and Met340. To test if Atg9 Phe17 is critical for binding to Atg23 we performed a pulldown assay with GST-Atg9 1-255 Phe17 mutated to Glu (Figure 3C). Replacing Phe17 with Glu resulted in a near complete loss of Atg23 binding demonstrating that Atg9 Phe17 is critical for binding to Atg23. Atg9 Phe17 was fully conserved across all 23 different yeast Atg9 sequences that we checked further highlighting the importance of this amino acid.

To test whether Atg23 cavity 1 is required for Atg9 binding we mutated Ala34, Leu323, Met340 and Ala377 and repeated our Atg23 pull-down assay. We also mutated Tyr16 which is predicted to face away from cavity 1 as we would predict that this mutation should have little effect on binding. Mutation of Atg23 Ala34, Leu323, Met340 or Ala377 led to significant reductions in Atg9 1-100 binding while mutation of Atg23 Tyr16 had no effect on Atg9 binding demonstrating that Atg9 is binding in cavity 1 on Atg23 (Figure 3F and 3G). To further explore the effect of these Atg23 mutations on the binding of Atg9 we also performed BLI with each of the different Atg23 mutants for Atg9 1-255 (Figure 3H and Table 2). Atg23 Leu323Arg, Met340Glu and Ala377Glu all had very little binding similar to the background signal and therefore a K_d_ could not be determined for these mutants. Atg23 Ala34Glu and Tyr16Glu had K_d_ values of 7.4 and 5.48 µM, respectively suggesting that these mutations had more modest effects on Atg9 binding.

### Atg23 interacts with the C-terminal tail of Atg9

The C-terminal intrinsically disordered region of Atg9 also has three regions of highly conserved amino acids. These are 870-888, 946-952, and 981-997 (Figure 3A). Given the high conservation in these regions, it is possible that these are also important protein-protein interaction regions. Therefore, we purified GST-Atg9 864-997 and tested its ability to bind to Atg23 using BLI (Figure 4A and S5A). Atg23 bound to Atg9 864-997 with a K_d_ of 0.46 ± 0.13 µM demonstrating that Atg23 can also bind to the C-terminus of Atg9 and that Atg23 has a higher affinity for the C-terminus than for the N-terminus (Table 2). We also utilized ITC to verify this binding interaction which revealed that the binding affinity of GST-Atg9 864-997 for Atg23 was 0.79 ± 0.12 µM with an n of 0.79 ± 0.2 which is very similar to the K_d_ measured by BLI and further confirms that Atg9 864-997 interacts with Atg23 (Figure S5B, S5C and Table 3).

**Figure 4.**
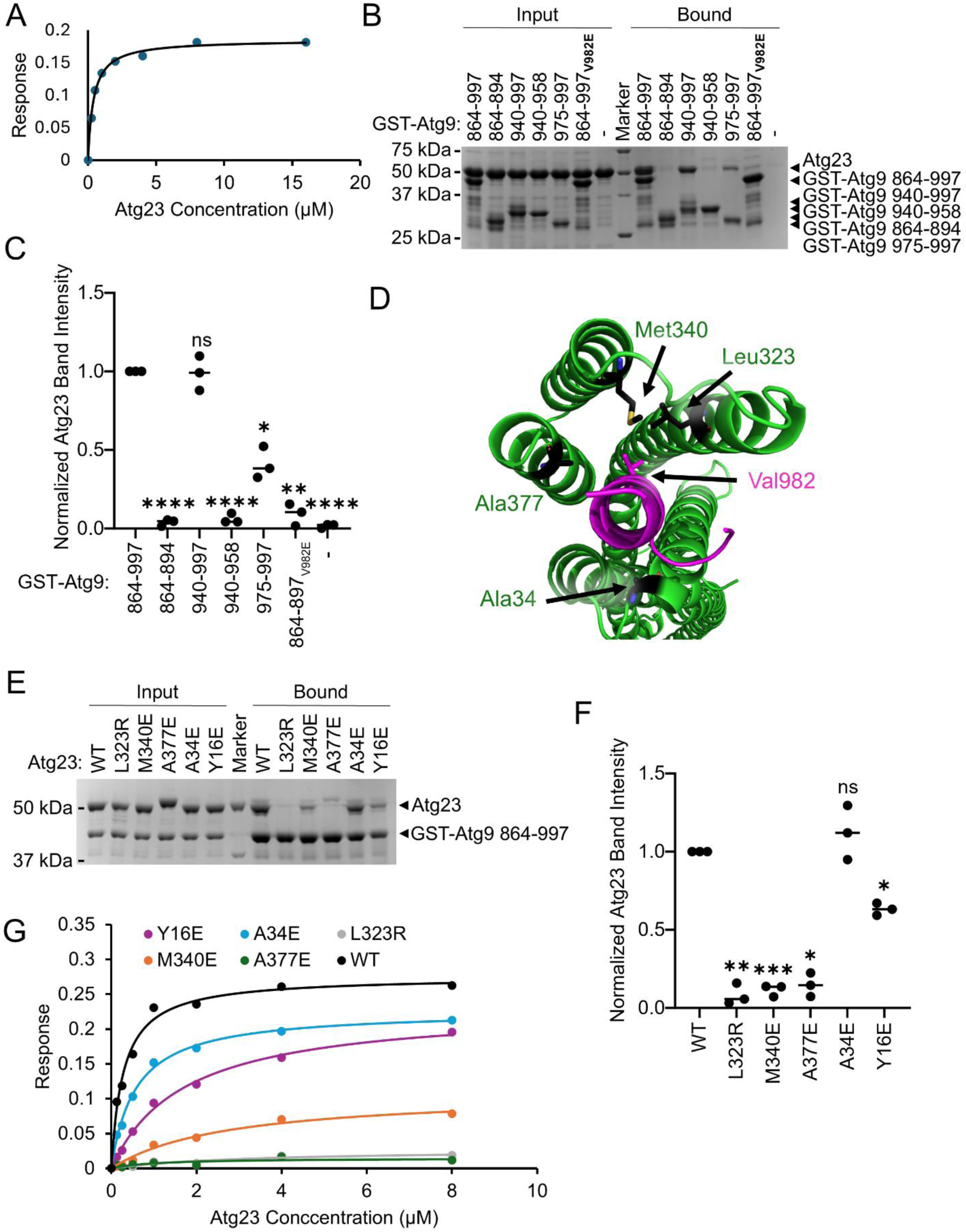
Atg23 interacts with the C-terminal region of Atg9. **A)** Steady state analysis of the BLI data for Atg9 849-997 and Atg23. Data is shown as dots and the fit to the data is shown as a line. **B)** SDS-PAGE gel for the GST-Atg9 C-terminal pulldowns of Atg23. The MW of each marker is shown on the left and the identity of each band is labeled on the right. **C)** Quantification of three independent repeats of B. Each replicate is shown as a dot and the average is shown as a line. **D)** Cartoon representation of the AlphaFold 3 model of Atg9 864-997 interaction with Atg23. Atg9 amino acids 977-997 are shown in magenta while the rest is hidden. Atg9 Val982 and Atg23 Ala34, Leu323, Met340 and Ala377 are all shown as stick representations and labeled. **E)** SDS-PAGE gel for the GST pulldown of Atg9 940-997 with different Atg23 point mutants. The MW of each marker is shown on the left and the identity of each band is labeled on the right. **F)** Quantification of three independent repeats of E shown as in C. **H)** Steady state analysis of the BLI data for Atg9 940-997 and different Atg23 point mutants. Data is shown as dots and the fit to the data is shown as a line. Statistical analysis in C and F was performed as a One-way ANOVA with data in C being compared to GST-Atg9 864-997 and data in F being compared to WT. **** P ≤ 0.0001, *** P ≤ 0.001, ** P ≤ 0.01, * P ≤ 0.05, ns, not significant.

To identify which region of the C-terminus of Atg9 is critical for Atg23 binding we generated a series of different GST tagged constructs of the C-terminal region based on the regions of sequence conservation. After purification of these constructs, we observed that they were more susceptible to proteolysis throughout the GST pulldown than the N-terminal region of Atg9. Despite this, we were still able to perform the GST pulldown assays as we did for Atg23 and the N-terminal region of Atg9. We found that Atg23 bound to GST-Atg9 940-997 similarly to GST-Atg9 846-997 suggesting that the main binding region is at the very C-terminus of Atg9 (Figure 4B and 4C). We next utilized AlphaFold 3 to predict the interaction region of Atg9 940-997. Atg9 940-997 docked into cavity 1 of Atg23 in a nearly identical location to the N-terminal region of Atg9 (Figure 4D), with a second surface-exposed binding site involving residues 946-952 of Atg9 contacting Tyr16 of Atg23. Atg9 982-997 formed a helix in the AlphaFold 3 prediction, and the helix was inserted into cavity 1 parallel to the helices from Atg23. This contrasts with the Atg9 N-terminal binding region which formed a helix that was perpendicular to the helices from Atg23 (Figure 3E). Atg9 Val982 is positioned in the hydrophobic portion of the cavity in the same location as Phe17 from the N-terminal Atg23 binding region. When Atg9 Val982 was mutated to Glu a near complete loss of Atg23 binding was observed for the 864-997 construct validating the importance of this amino acid in Atg23 binding (Figure 4B and 4C).

To determine whether Atg9 864-997 binds in cavity 1 on Atg23 we tested the binding of Atg23 Tyr16, Ala34, Leu323, Met340, and Ala377 mutants to GST-Atg9 864-997 (Figure 4E). Mutation of Leu323, Met340, and Ala377 showed a near complete loss in GST-Atg9 binding, demonstrating the importance of these amino acids for binding to the N-terminal region of Atg9 (Figure 4F). Intriguingly, mutation of Ala34 had essentially no effect on Atg9 864-997 binding which was not observed for the N-terminal region of Atg9 supporting that the binding sites of the N and C-terminal regions of Atg9 are largely overlapping but that they are not identical. To verify the effects of the Atg23 mutations we performed BLI with the different Atg23 mutants and Atg9 940-997. Atg23 bound to Atg9 940-997 with a K_d_ of 0.37 ± 0.13 µM which is very similar to the K_d_ for Atg23 binding to Atg9 864-997 further validating that this region is the main Atg23 interaction region (Figure 4G and Table 2). BLI data on Atg23 mutants revealed that Atg23 Leu323Arg, Met340Gluy and Ala377Glu all had very low binding similar to background and therefore a K_d_ could not be determined for these mutants. Atg23 Ala34Glu had a K_d_ of 0.57 µM indicating that this mutant had no effect on Atg9 869-997 binding which is consistent with the GST pulldown assay. Atg23 Tyr16Glu had a K_d_ of 1.72 µM for Atg9 869-997 indicating that this mutation resulted in a slight reduction in Atg9 binding.

## DISCUSSION

In this study, we report the structure of a monomeric form of Atg23, which we termed Atg23_ANNS_. Our experimental structure of Atg23_ANNS_ cannot be perfectly superimposed on the AlphaFold 3 model of dimeric Atg23 (Figure 2B and 2C). This occurs because of a difference in the angle of α7 pointing towards the dimerization region of Atg23 which results in a downward shift of α1, α2, α8, α9 and the second half of α7 of Atg23_ANNS_ relative to the AlphaFold 3 prediction (Figure 2C). This may suggest that the overall architecture of dimeric Atg23 is more curved than the AlphaFold 3 prediction or there is a flexible hinge in the middle of α7 and in between α2 and α3 that would allow α1, α2, α8, α9 and the second half of α7 to bend relative to the dimerization region of Atg23. If this is the case, then our crystal structure likely captured just a single state of this range of conformations.

In the structure of Atg23_ANNS_ we observed very weak helical density near where α6 is predicted to be located in the AlphaFold 3 model (Figure S3C). While we did not see any side chain density and therefore could not accurately assign these amino acids it is highly likely that this density corresponds to α6. Given the weak density for this helix in comparison to the remainder of the structure this region may be mobile and able to move relative to the rest of the structure. We also did not observe any electron density extending from α5 or α7 towards α6 which further supports the idea that this entire region is more dynamic than the rest of the structure. In the AlphaFold 3 model of dimeric Atg23 α6 forms an amphipathic helix where the hydrophobic surface of this helix points away from the rest of the structure and is accessible to solvent. Given that we previously demonstrated that Atg23 is membrane binding protein, it is possible that α6 is an important contributor to membrane binding and that the dynamics in this region are critical to enable membrane binding while Atg23 also interacts with Atg9 (55).

Another striking feature of the Atg23_ANNS_ structure is the large cavity formed by the helical bundle on the opposite end from where the dimer interface is located. This cavity is predominantly lined with conserved hydrophobic residues (Figure 2G). By combining different in vitro binding studies, we demonstrate that this cavity is the main binding site for both disordered termini of Atg9. Both termini contain a hydrophobic amino acid, Phe17 and Val982 which is located within a predicted alpha helix in Atg9 that docks into the center of the hydrophobic cavity in Atg23 and is essential for Atg23 binding. Both of these regions in Atg9 are distinct from the tandem PLF motifs (residues 163-165 and 187-189) that were previously shown to be involved in binding to the selective autophagy scaffolding protein Atg11 (28,29). Since the Atg23 and Atg11 binding motifs in Atg9 are so far apart it is possible that Atg9 may be able to bind to both Atg23 and Atg11 simultaneously, but this will need to be addressed in future work. In addition, the N-terminal Atg23 binding region in Atg9 includes Ser19 which was previously identified as a phosphorylation site (59). This suggests that the interaction between Atg23 and Atg9 could be partially regulated by phosphorylation in the N-terminal region of Atg9. A second phosphorylation site, Ser948, has also been identified in Atg9 (59). This phosphorylation site is near the C-terminal Atg23 binding motif but is not directly overlapping and as such it is less clear whether phosphorylation at this site could regulate the interaction between Atg23 and Atg9.

We determined that the K_d_ for the interaction between Atg23 and Atg9 is 5 µM for the N-terminus and 0.78 µM for the C-terminus by ITC and 1.5 µM and 0.45 µM, respectively, by BLI. The low affinity interaction between Atg23 and Atg9 is consistent with several other autophagy protein interactions including, the interaction between Atg11 and Atg9 which was reported at 1.09 µM (59). The abundance of Atg23 and Atg9 are nearly stoichiometric, with roughly 1000 copies per yeast cell equating to a cellular concentration of 50 nM intracellular concentration (60). With such a low concentration in the cell a 1 to 1 interaction between Atg23 and Atg9 would not be predicted to be stable. However, Atg23 is a dimer and Atg9 is a trimer in solution. In addition, since Atg23 is able to interact with both the N- and C- termini of Atg9 it is possible that the interaction between Atg23 and Atg9 results in an avidity driven clustering effect which has been observed for many autophagy proteins (61). This is in line with both the punctate localization for Atg23 and Atg9 and is a recurring theme within autophagy protein interactions.

## EXPERIMENTAL PROCEDURES

### Reagents

The following reagents and consumables were used in thus study: Octet HIS1K Biosensors (Sartorius 18-5120), sodium chloride (Therm. Sci. 447300010), Tris Buffer (Invitrogen 15504-020), imidazole (Therm. Sci. J17525-A1), Phenylmethanesulfonyl fluoride (PMSF) (VWR 0754-25), cOmplete EDTA-free protease inhibitor tablets (Roche 11836170001) TALON affinity resin (Takara 635504), Morpheus I (Molecular Dimensions MD1-46), Morpheus II (Molecular Dimensions MD1-92), Precipitant Mix 7 (Molecular Dimensions MD2-100-241), Monosaccharides II (Molecular Dimensions MD2-100-236), Buffer System 4 (Molecular Dimensions MD2-250-243), glutathione Sepharose 4B (GE Healthcare 17-0756-05), glutathione (Affymetrix 16315), 37% hydrochloric acid (EMD Millipore, HX0603-4), Sodium Dodecyl Sulfate (SDS)(Invitrogen 15525017), glycine (Invitrogen 15527013), 40% acrylamide/bis-acrylamide (37.5:1, Bio-Rad 1610148), ammonium persulfate (Bio-Rad 1610700), tetramethylethylenediamine (Bio-Rad 1610800), methanol (Fisher A412-4), acetic acid (Supelco AX0073-9), Coomassie Brilliant Blue G-250 (VWR M140) LB Broth Miller (Fisher BP1426-2), glucose (Sigma G6152), Isopropyl β-D-thiogalatopyranoside (IPTG)(IBI Scientific IB02125), selenomethionine (Molecular Dimensions MD12-503B), potassium phosphate monobasic (Sigma P5655-500), sodium phosphate dibasic (Fisher S374-500), ammonium chloride (Fisher A661-500), BME vitamins (Sigma B6891), lysine (TCI L0129), phenylalanine (TCI P0134), threonine (TCI T0230), leucine (Alfa Aesar A12311), valine (Alfa Aesar A12720), isoleucine (TCI I0181), agar (Therm. Sci. A10752.36), chloramphenicol (Fisher BP904-100), ampicillin (Fisher BP1760-25), kanamycin (Gibco 11815-032), Q5 2X master mix (M0492S), KLD enzyme mix (NEB M0554S), 2x Gibson Assembly master mix (NEB E2611S), T4 ligase (NEB M0202S); BamHI-HF, SalI-HF, NcoI-HF, NotI-HF, DpnI (NEB R3136L, R3138S, R3193S, R3189S, R0176S); 2x GoTaq master mix (Promega M7122), NEB5 alpha (NEB C2987H), Rosetta2 (DE3) (Novagen 71397), QiaQuick PCR Purification Kit (Qiagen 28104), and QiaPrep Spin Miniprep Kit (Qiagen 27104).

### Plasmids and cloning

A complete list of the plasmids used in this manuscript are available in supplemental table 1. Atg23 constructs (1-381 and full-length) were PCR amplified, column-purified, digested, heat inactivated and ligated into the digested and column-purified pHIS-parallel2 vector between the BamHI and NotI sites, after which deletion (ANNS mutant of 1-381) and mutations (Leu323Arg, Met340Glu, Ala377Glu, Ala34Glu, Tyr16Glu) were introduced using the Q5 site-directed mutagenesis protocol (62). Atg9 constructures were generated using a similar approach to Atg23 expect that they were cloned into the pGST2 vector containing a 10xHis tag prior to the GST tag and TEV cleavage site (62). Atg9 1-100 and 40-235 were ligated into the BamHI and NotI sites while Atg9 1-255, 864-997, 864-894 and 940-997 were ligated into the NcoI and SalI sites. Further deletions (940–958, 975–997) and mutations (Phe17Glu, Val982Glu) were introduced by Q5 site directed mutagenesis. GST-Atg9 12-24 was generated by insertional mutagenesis into the pET His6 GST TEV LIC cloning vector (1G), which was a gift from Scott Gradia (Addgene plasmid # 29655) using Q5 site directed mutagenesis. 6xHis-Atg9-1-255-TEV-MBP constructs was generated by Gibson cloning between the His-tag and MBP of the pET His6 MBP TEV LIC cloning vector (1M), which was a gift from Scott Gradia (Addgene plasmid # 29656), and additional constructs were generated via Q5 site directed mutagenesis. Neb5 alpha cells were transformed according to the NEB protocol, screened with colony PCR, miniprepped and verified by Nanopore plasmid sequencing at the Genomics and Molecular Biology Shared Resources Core at Dartmouth.

### Protein expression

2.8L Fernbach flasks were used for expressing protein in Rosetta2 (DE3) bacteria. For all Atg23 constructs (monomeric Atg23_ANNS_, full-length, and its mutants), Rosetta2 (DE3) chemically competent cells were transformed with 100 ng of the corresponding plasmid, plated on chloramphenicol and ampicillin LB Miller agar, grown overnight at 37°C and directly transferred to 1L of LB with 34 mg/L chloramphenicol, 100 mg/L ampicillin, grown with shaking at 37°C until O.D. at 600 nm reached 0.5, temperature was set to lower to 18℃ and after 30 minutes of cooling, IPTG was added to 0.2 mM final, with expression proceeding overnight at 18°C.

For selenomethionine labeling, freshly transformed Rosetta2 (DE3) cells on a plate were directly transferred to 1L of M9 minimal media (5.8 g sodium phosphate dibasic, 3.0 g potassium phosphate monobasic, 0.5 g sodium chloride, 1.0 g ammonium chloride, 0.5 g magnesium sulfate, 4 g glucose, 10 mL 100x BME vitamins, 2 mL trace metal solution (5 g FeCl_2_*2H_2_O, 184 mg CaCl_2_*2H_2_O, 64 mg H_3_BO_3_, 18 mg CoCl_2_*6H_2_O, 4 mg CuCl_2_*2H_2_O, 340 mg ZnCl_2_, 40 mg Na_2_MoO_4_*2H_2_O, 3 mL 12 M HCl per 1L) filter-sterilized), grown at 37℃ until O.D. at 600 nm reached 0.5, cooled for 30 minutes to 18°C, selenomethionine (60 mg), phenylalanine (100 mg), lysine (100 mg), threonine (100 mg), leucine (50 mg), valine (50 mg), isoleucine (50 mg) were added to supplement the selenomethionine and suppress *de novo* amino acid synthesis from aspartate, and cultures were induced with 1 mM final IPTG, growing overnight at 18℃.

For expressing GST-Atg9 fusions and Atg9-TEV-MBP fusions, freshly transformed Rosetta2 (DE3) were transferred to 1L of LB with 34 mg/L chloramphenicol, 100 mg/L ampicillin, 2 g/L glucose to suppress initial expression and grown at 37°C to O.D. 1 at 600 nm. IPTG was added to 0.2 mM final and expressed further at 37°C for 3 hours to reduce the proteolytic degradation.

### Protein purification

After growth in the presence of IPTG, cells were harvested at 4400 g, 4°C for 25 minutes in a JS-4.2 rotor, media decanted and the cells resuspended in cold IMAC binding buffer (500 mM NaCl, 20 mM Tris pH8 for Atg23 or 20 mM sodium phosphate pH 7 for Atg9) at a 1:3 cell pellet:buffer ratio and protease inhibitors were added (1/3 cOmplete EDTA-free tablet per liter, 1 mM PMSF final concentration). Cells were lysed by 3 passes at 18,000 psi through the LM10 microfluidic processor with a cooling coil submerged into ice-water bath, the lysate was clarified at 55,000 g, 4℃ for 20 minutes in JA-25.50 rotor and applied on 4 mL of TALON resin packed into 16 mm diameter column, the column was washed with 16 mL of IMAC binding buffer, then with 16 mL of 150 mM NaCl, 20 mM Tris pH8 (TBS) supplemented with 2.5 mM imidazole, and eluted in 4x 4 mL fractions of TBS supplemented with 150 mM imidazole, discarding the first 1/3 column volume as the dead volume. For Atg23 and the Atg9-TEV-MBP, 2 mM final DTT, EDTA were added and the tags were cleaved with TEV protease at a 1:100 molar ratio for 16 hours at 4°C. For Atg9 1-255 – MBP, the protein was diluted 3 times with water, applied on 5 mL of HiTrap Q HP column in TBS and eluted with 50-1000 mM NaCl, 20 mM Tris pH8 gradient; MBP elutes at 150 mM NaCl while Atg9 at 500 mM NaCl. After tag cleavage or directly for GST-Atg9 fusions, protein was further purified by SEC using Superdex200 16/600 (or Superdex75 16/600 for monomeric Atg23 and Atg9, 1-48) column equilibrated in 150 mM NaCl, 20 mM Tris pH8, 0.2 mM Tris(2-carboxyethyl) phosphine (TCEP). Peak fractions were pooled, concentrated and frozen in liquid nitrogen.

### Protein crystallization and structure determination

Monomeric Atg23_ANNS_ at 8 mg/mL in TBS with 0.2 mM TCEP was screened using sparse-matrix Morpheus I and II screens, setting 200 nL with 200 nL and 133 nL with 267 nL protein plus reservoir drops in 96-well UVXPO MRC plates using the NT8 drop setter (63,64). Needle cluster crystals grew in the MPM6, BS6, polyamines II condition; numerous other conditions were giving sea urchin-like structures, but the best, longest and well-formed crystals grew from MPM7 (10% PEG-8000, 20% 1,5-pentanediol), BS4 (0.07M MOPSO, 0.03M Bis-Tris), Monosaccharides II (20 mM each of xylitol, myo-inositol, D-(-)-fructose, L-rhamnose, D-sorbitol) condition at 1:2 protein:reservoir ratio. They could be reproduced well in 24-well plates with fine-screening showing that adjusting precipitant concentration to 8.8% PEG-8000, 17.6% 1,5-pentanediol, while keeping other components identical, gave the largest crystals with seeding at room temperature after 1 day. Adding TCEP to 2 mM helped to eliminate the appearance of denatured protein skin on the drops. Selenomethionine-labeled crystals were frozen directly in liquid nitrogen as the condition is cryo-protected and a vector data collection was performed along the long rod-like crystals on the selenium edge. Raw images were subjected to spot finding and indexing using XDS, excluding the start and end portions of the vector data collection because of lower data quality (65–69). Reflections were scaled using AIMLESS, after which the Phaser MR-SAD module in Phenix was used for phasing, with a truncated Atg23_ANNS_ AF2 search model (only residues 1-65, 288-381 of the full-length Atg23 were included for MR) (70–74). Autobuild module in Phenix was used to build an initial model, which was modified in Coot and refined in Phenix (75–77). The structure of Atg23_ANNS_ has been deposited in the PDB with ID: 9ZM0.

### GST pulldowns

Atg23 constructs were diluted with 150 mM NaCl, 20 mM Tris pH8 (TBS), after which the indicated GST-Atg9 construct was added, so that the final concentrations of both proteins would be 5 µM for pulldowns with Atg9 N-terminal constructs or 2 µM for the C-terminal Atg9 constructs, 200 µL final volume, 25 µL sample was then taken and 25 µL of 50% glutathione agarose 4B slurry equilibrated in TBS was added and incubated at 4°C for 15 minutes with inverting. Beads were spun down at 1000 g, 1 minute, room temperature; the supernatant was removed, and the beads were washed with 500 µL of TBS for 5 minutes at room temperature, spun down at 1000 g, 1 minute, room temperature.; the supernatant removed, and the beads were eluted with 30 µL of TBS with 10 mM neutralized glutathione. 10 µL of sample was loaded on a 15-well 10% acrylamide Tris-glycine gel, with 192 mM glycine, 25 mM Tris, 0.1% SDS running buffer, stained in 20% methanol, 10% acetic acid, 0.1% Coomassie G-250 for 30 minutes, destained in 10% methanol, 5% acetic acid for 3 hours and further destained in water. Bands were quantified in ImageLab 3.1 (BioRad) using the volume tools or ImageJ, taking the lowest intensity pixel as the background signal (78). For mutant pulldowns, signal was calculated as the ratio of Atg23 to Atg9 band intensity, as when no Atg23 binds more GST-Atg9 fusion is able to bind the beads, potentially because of more available space without Atg23 on the beads; for truncations, raw Atg23 band intensity was used. Values were normalized to that of wild-type or full-length tail construct, and statistical analysis was performed in GraphPad Prism using the ANOVA one-way analysis.

### BLI

HIS1K chips were activated by submersion for 3 minutes in TBS with 0.1% Tween-20. After this, TBS with 0.02% Tween-20 was used as buffer for all samples. The Octet BLI system was programmed to measure baseline in buffer for 60 seconds, load His-tagged Atg9 for 120-200 seconds, measure the baseline in buffer again for 120 seconds, bind Atg23 for 200 seconds and measure dissociation in buffer for 200 seconds. Tips were then regenerated by 3 sequential 5-second dips into 10 mM glycine pH 3 and buffer. Temperature was set to 30°C, shake speed was 300 rpm and the data recorded every 0.2 seconds.

6xHis-Atg9 1-255 was used at 2 µM for loading, all other 10xHis-GST-Atg9 constructs were loaded at 0.5 µM. Atg23 was used at 0, 0.25, 0.5, 1, 2, 4, 8 and 16 µM for binding; mutants were measured at 0, 0.125, 0.25, 0.5, 1, 2, 4, 8 µM.

For data analysis, 20 points (2 seconds of real time window) were averaged 5 seconds before binding start to give the initial signal, then 20 points were averaged 5 seconds before binding completion to give the final signal. All sensorgrams were aligned to the 5 second before binding start point for display. The amount of Atg23 bound was calculated as the difference between these two values, adjusted for some background dissociation of the His-Atg9 construct by adding the difference from the 0 µM Atg23 well. Non-linear curve fit was then performed to fit the binding with an independent model and dissociation constants were averaged for three replicates where applicable, the error being the standard deviation of the replicates.

### ITC

Frozen Atg23 aliquot at 220 µM and Atg9 1-255 or Atg9 864-997 at 22 µM were thawed on ice, dialyzed together against 150 mM NaCl, 20 mM Tris pH8, 0.2 mM TCEP at 4°C for 4 hours and degassed at room temperature for 30 minutes under vacuum. TA instruments Nano ITC with 1 mL cell and 250 µL syringe was thoroughly washed with water, dried by passing air through the cell and the syringe, 1.2 mL of Atg9 was loaded into the cell avoiding bubbles, while 210 µL of Atg23 was loaded into syringe, leaving a small bubble above the solution. Temperature of the cell was set to 20°C and stir speed to 150 rpm. System was equilibrated until a stable baseline was established and twenty 10µL injections were performed with 200 s intervals between injections. Baseline was automatically determined and data fit with independent binding model in NanoAnalyze. Dissociation constants from 3 (N-terminal) or 4 (C-terminal) runs were averaged, and the error is presented as standard deviation

## Data Availability

The crystal structure of Atg23_ANNS_ is available on the PDB (ID: 9ZM0).

## Acknowledgements

We thank the beamline staff at AMX and FMX for their help throughout this project. We also thank Dr. Grace Ge and Dr. Margaret E. Ackerman for the help with bio-layer interferometry and Dr. Zdenek Svindrych, along with all BioMT staff for the help with high-throughput crystallization and isothermal titration calorimetry.

## Author contributions

**ZB:** conceptualization, data curation, formal analysis, investigation, methodology, validation, visualization, writing – original draft, writing – review & editing

**KAL:** conceptualization, methodology, writing – review & editing

**MJR:** conceptualization, funding acquisition, project administration, supervision, writing – original draft, writing – review & editing

## References

1. Wen, X., and Klionsky, D. J. (2016) An overview of macroautophagy in yeast. J Mol Biol 428, 1681–1699

2. Lamark, T., and Johansen, T. (2021) Mechanisms of Selective Autophagy. Annu Rev Cell Dev Biol 37, 143–169

3. Morishita, H., and Mizushima, N. (2019) Diverse Cellular Roles of Autophagy. Annu Rev Cell Dev Biol 35, 453–475

4. Dikic, I., and Elazar, Z. (2018) Mechanism and medical implications of mammalian autophagy. Nat Rev Mol Cell Biol 19, 349–364

5. Klionsky, D. J., Petroni, G., Amaravadi, R. K., Baehrecke, E. H., Ballabio, A., Boya, P., Bravo-San Pedro, J. M., Cadwell, K., Cecconi, F., Choi, A. M. K., Choi, M. E., Chu, C. T., Codogno, P., Colombo, M. I., Cuervo, A. M., Deretic, V., Dikic, I., Elazar, Z., Eskelinen, E. L., Fimia, G. M., Gewirtz, D. A., Green, D. R., Hansen, M., Jaattela, M., Johansen, T., Juhasz, G., Karantza, V., Kraft, C., Kroemer, G., Ktistakis, N. T., Kumar, S., Lopez-Otin, C., Macleod, K. F., Madeo, F., Martinez, J., Melendez, A., Mizushima, N., Munz, C., Penninger, J. M., Perera, R. M., Piacentini, M., Reggiori, F., Rubinsztein, D. C., Ryan, K. M., Sadoshima, J., Santambrogio, L., Scorrano, L., Simon, H. U., Simon, A. K., Simonsen, A., Stolz, A., Tavernarakis, N., Tooze, S. A., Yoshimori, T., Yuan, J., Yue, Z., Zhong, Q., Galluzzi, L., and Pietrocola, F. (2021) Autophagy in major human diseases. EMBO J 40, e108863

6. Qu, X., Yu, J., Bhagat, G., Furuya, N., Hibshoosh, H., Troxel, A., Rosen, J., Eskelinen, E. L., Mizushima, N., Ohsumi, Y., Cattoretti, G., and Levine, B. (2003) Promotion of tumorigenesis by heterozygous disruption of the beclin 1 autophagy gene. J Clin Invest 112, 1809–1820

7. Wang, L., Klionsky, D. J., and Shen, H. M. (2023) The emerging mechanisms and functions of microautophagy. Nat Rev Mol Cell Biol 24, 186–203

8. Kaushik, S., and Cuervo, A. M. (2018) The coming of age of chaperone-mediated autophagy. Nat Rev Mol Cell Biol 19, 365–381

9. Nakatogawa, H. (2020) Mechanisms governing autophagosome biogenesis. Nat Rev Mol Cell Biol 21, 439–458

10. Yamamoto, H., Kakuta, S., Watanabe, T. M., Kitamura, A., Sekito, T., Kondo-Kakuta, C., Ichikawa, R., Kinjo, M., and Ohsumi, Y. (2012) Atg9 vesicles are an important membrane source during early steps of autophagosome formation. J Cell Biol 198, 219–233

11. Mari, M., Griffith, J., Rieter, E., Krishnappa, L., Klionsky, D. J., and Reggiori, F. (2010) An Atg9-containing compartment that functions in the early steps of autophagosome biogenesis. J Cell Biol 190, 1005–1022

12. Olivas, T. J., Wu, Y., Yu, S., Luan, L., Choi, P., Guinn, E. D., Nag, S., De Camilli, P. V., Gupta, K., and Melia, T. J. (2023) ATG9 vesicles comprise the seed membrane of mammalian autophagosomes. J Cell Biol 222

13. Schutter, M., Giavalisco, P., Brodesser, S., and Graef, M. (2020) Local Fatty Acid Channeling into Phospholipid Synthesis Drives Phagophore Expansion during Autophagy. Cell 180, 135–149 e114

14. Dabrowski, R., Tulli, S., and Graef, M. (2023) Parallel phospholipid transfer by Vps13 and Atg2 determines autophagosome biogenesis dynamics. J Cell Biol 222

15. Osawa, T., Kotani, T., Kawaoka, T., Hirata, E., Suzuki, K., Nakatogawa, H., Ohsumi, Y., and Noda, N. N. (2019) Atg2 mediates direct lipid transfer between membranes for autophagosome formation. Nat Struct Mol Biol 26, 281–288

16. Wang, Y., Dahmane, S., Ti, R., Mai, X., Zhu, L., Carlson, L. A., and Stjepanovic, G. (2025) Structural basis for lipid transfer by the ATG2A-ATG9A complex. Nat Struct Mol Biol 32, 35–47

17. Valverde, D. P., Yu, S., Boggavarapu, V., Kumar, N., Lees, J. A., Walz, T., Reinisch, K. M., and Melia, T. J. (2019) ATG2 transports lipids to promote autophagosome biogenesis. J Cell Biol 218, 1787–1798

18. Takahashi, Y., He, H., Tang, Z., Hattori, T., Liu, Y., Young, M. M., Serfass, J. M., Chen, L., Gebru, M., Chen, C., Wills, C. A., Atkinson, J. M., Chen, H., Abraham, T., and Wang, H. G. (2018) An autophagy assay reveals the ESCRT-III component CHMP2A as a regulator of phagophore closure. Nat Commun 9, 2855

19. Itakura, E., Kishi-Itakura, C., and Mizushima, N. (2012) The hairpin-type tail-anchored SNARE syntaxin 17 targets to autophagosomes for fusion with endosomes/lysosomes. Cell 151, 1256–1269

20. Kirisako, T., Baba, M., Ishihara, N., Miyazawa, K., Ohsumi, M., Yoshimori, T., Noda, T., and Ohsumi, Y. (1999) Formation process of autophagosome is traced with Apg8/Aut7p in yeast. Journal of Cell Biology 147, 435–446

21. May, A. I., Prescott, M., and Ohsumi, Y. (2020) Autophagy facilitates adaptation of budding yeast to respiratory growth by recycling serine for one-carbon metabolism. Nat Commun 11, 5052

22. Guiboileau, A., Yoshimoto, K., Soulay, F., Bataillé, M. P., Avice, J. C., and Masclaux-Daubresse, C. (2012) Autophagy machinery controls nitrogen remobilization at the whole-plant level under both limiting and ample nitrate conditions in Arabidopsis. New Phytol 194, 732–740

23. Feng, Y., He, D., Yao, Z., and Klionsky, D. J. (2014) The machinery of macroautophagy. Cell Res 24, 24–41

24. Guardia, C. M., Tan, X. F., Lian, T., Rana, M. S., Zhou, W., Christenson, E. T., Lowry, A. J., Faraldo-Gomez, J. D., Bonifacino, J. S., Jiang, J., and Banerjee, A. (2020) Structure of Human ATG9A, the Only Transmembrane Protein of the Core Autophagy Machinery. Cell Rep 31, 107837

25. Lang, T., Reiche, S., Straub, M., Bredschneider, M., and Thumm, M. (2000) Autophagy and the cvt pathway both depend on AUT9. J Bacteriol 182, 2125–2133

26. Ohashi, Y., and Munro, S. (2010) Membrane delivery to the yeast autophagosome from the Golgi-endosomal system. Mol Biol Cell 21, 3998–4008

27. Matscheko, N., Mayrhofer, P., Rao, Y., Beier, V., and Wollert, T. (2019) Atg11 tethers Atg9 vesicles to initiate selective autophagy. PLoS Biol 17, e3000377

28. He, C., Song, H., Yorimitsu, T., Monastyrska, I., Yen, W. L., Legakis, J. E., and Klionsky, D. J. (2006) Recruitment of Atg9 to the preautophagosomal structure by Atg11 is essential for selective autophagy in budding yeast. J Cell Biol 175, 925–935

29. Coudevylle, N., Banas, B., Baumann, V., Schuschnig, M., Zawadzka-Kazimierczuk, A., Kozminski, W., and Martens, S. (2022) Mechanism of Atg9 recruitment by Atg11 in the cytoplasm-to-vacuole targeting pathway. J Biol Chem 298, 101573

30. Sekito, T., Kawamata, T., Ichikawa, R., Suzuki, K., and Ohsumi, Y. (2009) Atg17 recruits Atg9 to organize the pre-autophagosomal structure. Genes Cells 14, 525–538

31. Yorimitsu, T., and Klionsky, D. J. (2005) Atg11 links cargo to the vesicle-forming machinery in the cytoplasm to vacuole targeting pathway. Mol Biol Cell 16, 1593–1605

32. Maeda, S., Yamamoto, H., Kinch, L. N., Garza, C. M., Takahashi, S., Otomo, C., Grishin, N. V., Forli, S., Mizushima, N., and Otomo, T. (2020) Structure, lipid scrambling activity and role in autophagosome formation of ATG9A. Nature Structural & Molecular Biology 27, 1194–U1246

33. Matoba, K., Kotani, T., Tsutsumi, A., Tsuji, T., Mori, T., Noshiro, D., Sugita, Y., Nomura, N., Iwata, S., Ohsumi, Y., Fujimoto, T., Nakatogawa, H., Kikkawa, M., and Noda, N. N. (2020) Atg9 is a lipid scramblase that mediates autophagosomal membrane expansion. Nature Structural & Molecular Biology 27, 1185–U1224

34. Arlt, H., Raman, B., Filali-Mouncef, Y., Hu, Y., Leytens, A., Hardenberg, R., Guimaraes, R., Kriegenburg, F., Mari, M., Smaczynska-de, R., II, Ayscough, K. R., Dengjel, J., Ungermann, C., and Reggiori, F. (2023) The dynamin Vps1 mediates Atg9 transport to the sites of autophagosome formation. J Biol Chem 299, 104712

35. Darsow, T., Rieder, S. E., and Emr, S. D. (1997) A multispecificity syntaxin homologue, Vam3p, essential for autophagic and biosynthetic protein transport to the vacuole. J Cell Biol 138, 517–529

36. Geng, J. F., Nair, U., Yasumura-Yorimitsu, K., and Klionsky, D. J. (2010) Post-Golgi Sec Proteins Are Required for Autophagy in. Molecular Biology of the Cell 21, 2257–2269

37. Ishihara, N., Hamasaki, M., Yokota, S., Suzuki, K., Kamada, Y., Kihara, A., Yoshimori, T., Noda, T., and Ohsumi, Y. (2001) Autophagosome requires specific early Sec proteins for its formation and NSF/SNARE for vacuolar fusion. Mol Biol Cell 12, 3690–3702

38. Kakuta, S., Yamamoto, H., Negishi, L., Kondo-Kakuta, C., Hayashi, N., and Ohsumi, Y. (2012) Atg9 vesicles recruit vesicle-tethering proteins Trs85 and Ypt1 to the autophagosome formation site. J Biol Chem 287, 44261–44269

39. Kriegenburg, F., Huiting, W., van Buuren-Broek, F., Zwilling, E., Hardenberg, R., Mari, M., Kraft, C., and Reggiori, F. (2022) The lipid flippase Drs2 regulates anterograde transport of Atg9 during autophagy. Autophagy Rep 1, 345–367

40. Lipatova, Z., Belogortseva, N., Zhang, X. Q., Kim, J., Taussig, D., and Segev, N. (2012) Regulation of selective autophagy onset by a Ypt/Rab GTPase module. Proc Natl Acad Sci U S A 109, 6981–6986

41. Lynch-Day, M. A., Bhandari, D., Menon, S., Huang, J., Cai, H., Bartholomew, C. R., Brumell, J. H., Ferro-Novick, S., and Klionsky, D. J. (2010) Trs85 directs a Ypt1 GEF, TRAPPIII, to the phagophore to promote autophagy. Proc Natl Acad Sci U S A 107, 7811–7816

42. Reggiori, F., Wang, C. W., Nair, U., Shintani, T., Abeliovich, H., and Klionsky, D. J. (2004) Early stages of the secretory pathway, but not endosomes, are required for Cvt vesicle and autophagosome assembly in Saccharomyces cerevisiae. Mol Biol Cell 15, 2189–2204

43. Reggiori, F., Wang, C. W., Stromhaug, P. E., Shintani, T., and Klionsky, D. J. (2003) Vps51 is part of the yeast Vps fifty-three tethering complex essential for retrograde traffic from the early endosome and Cvt vesicle completion. Journal of Biological Chemistry 278, 5009–5020

44. Shirahama-Noda, K., Kira, S., Yoshimori, T., and Noda, T. (2013) TRAPPIII is responsible for vesicular transport from early endosomes to Golgi, facilitating Atg9 cycling in autophagy. Journal of Cell Science 126, 4963–4973

45. von Mollard, G. F., and Stevens, T. H. (1999) The v-SNARE Vti1p is required for multiple membrane transport pathways to the vacuole. Molecular Biology of the Cell 10, 1719–1732

46. Wang, I. H., Chen, Y. J., Hsu, J. W., and Lee, F. J. (2017) The Arl3 and Arl1 GTPases co-operate with Cog8 to regulate selective autophagy via Atg9 trafficking. Traffic 18, 580–589

47. Yang, S., and Rosenwald, A. G. (2016) Autophagy in Saccharomyces cerevisiae requires the monomeric GTP-binding proteins, Arl1 and Ypt6. Autophagy 12, 1721–1737

48. Yen, W. L., Shintani, T., Nair, U., Cao, Y., Richardson, B. C., Li, Z. J., Hughson, F. M., Baba, M., and Klionsky, D. J. (2010) The conserved oligomeric Golgi complex is involved in double-membrane vesicle formation during autophagy. Journal of Cell Biology 188, 101–114

49. Zou, S. S., Sun, D., and Liang, Y. H. (2017) The Roles of the SNARE Protein Sed5 in Autophagy in. Mol Cells 40, 643–654

50. Tucker, K. A., Reggiori, F., Dunn, W. A., and Klionsky, D. J. (2003) Atg23 is essential for the cytoplasm to vacuole targeting pathway and efficient autophagy but not pexophagy. Journal of Biological Chemistry 278, 48445–48452

51. Reggiori, F., Tucker, K. A., Stromhaug, P. E., and Klionsky, D. J. (2004) The Atg1-Atg13 complex regulates Atg9 and Atg23 retrieval transport from the pre-autophagosomal structure. Developmental Cell 6, 79–90

52. Yen, W. L., Legakis, J. E., Nair, U., and Klionsky, D. J. (2007) Atg27 is required for autophagy-dependent cycling of Atg9. Molecular Biology of the Cell 18, 581–593

53. Legakis, J. E., Yen, W. L., and Klionsky, D. J. (2007) A cycling protein complex required for selective autophagy. Autophagy 3, 422–432

54. Backues, S. K., Orban, D. P., Bernard, A., Singh, K., Cao, Y., and Klionsky, D. J. (2015) Atg23 and Atg27 act at the early stages of Atg9 trafficking in S. cerevisiae. Traffic 16, 172–190

55. Hawkins, W. D., Leary, K. A., Andhare, D., Popelka, H., Klionsky, D. J., and Ragusa, M. J. (2022) Dimerization-dependent membrane tethering by Atg23 is essential for yeast autophagy. Cell Rep 39, 110702

56. Abramson, J., Adler, J., Dunger, J., Evans, R., Green, T., Pritzel, A., Ronneberger, O., Willmore, L., Ballard, A. J., Bambrick, J., Bodenstein, S. W., Evans, D. A., Hung, C. C., O’Neill, M., Reiman, D., Tunyasuvunakool, K., Wu, Z., Zemgulyte, A., Arvaniti, E., Beattie, C., Bertolli, O., Bridgland, A., Cherepanov, A., Congreve, M., Cowen-Rivers, A. I., Cowie, A., Figurnov, M., Fuchs, F. B., Gladman, H., Jain, R., Khan, Y. A., Low, C. M. R., Perlin, K., Potapenko, A., Savy, P., Singh, S., Stecula, A., Thillaisundaram, A., Tong, C., Yakneen, S., Zhong, E. D., Zielinski, M., Zidek, A., Bapst, V., Kohli, P., Jaderberg, M., Hassabis, D., and Jumper, J. M. (2024) Accurate structure prediction of biomolecular interactions with AlphaFold 3. Nature 630, 493–500

57. van Kempen, M., Kim, S. S., Tumescheit, C., Mirdita, M., Lee, J., Gilchrist, C. L. M., Soding, J., and Steinegger, M. (2024) Fast and accurate protein structure search with Foldseek. Nat Biotechnol 42, 243–246

58. Xu, Y., Wang, S., Hu, Q., Gao, S., Ma, X., Zhang, W., Shen, Y., Chen, F., Lai, L., and Pei, J. (2018) CavityPlus: a web server for protein cavity detection with pharmacophore modelling, allosteric site identification and covalent ligand binding ability prediction. Nucleic Acids Res 46, W374–W379

59. Papinski, D., Schuschnig, M., Reiter, W., Wilhelm, L., Barnes, C. A., Maiolica, A., Hansmann, I., Pfaffenwimmer, T., Kijanska, M., Stoffel, I., Lee, S. S., Brezovich, A., Lou, J. H., Turk, B. E., Aebersold, R., Ammerer, G., Peter, M., and Kraft, C. (2014) Early steps in autophagy depend on direct phosphorylation of Atg9 by the Atg1 kinase. Mol Cell 53, 471–483

60. Geng, J. F., Baba, M., Nair, U., and Klionsky, D. J. (2008) Quantitative analysis of autophagy-related protein stoichiometry by fluorescence microscopy. Journal of Cell Biology 182, 129–140

61. Hollenstein, D. M., Licheva, M., Konradi, N., Schweida, D., Mancilla, H., Mari, M., Reggiori, F., and Kraft, C. (2021) Spatial control of avidity regulates initiation and progression of selective autophagy. Nature Communications 12

62. Sheffield, P., Garrard, S., and Derewenda, Z. (1999) Overcoming expression and purification problems of RhoGDI using a family of “parallel” expression vectors. Protein Expr Purif 15, 34–39

63. Gorrec, F. (2009) The MORPHEUS protein crystallization screen. J Appl Crystallogr 42, 1035–1042

64. Gorrec, F. (2015) The MORPHEUS II protein crystallization screen. Acta Crystallogr F Struct Biol Commun 71, 831–837

65. Kabsch, W. (1988) Evaluation of Single-Crystal X-Ray-Diffraction Data from a Position-Sensitive Detector. Journal of Applied Crystallography 21, 916–924

66. Kabsch, W. (1988) Automatic-Indexing of Rotation Diffraction Patterns. Journal of Applied Crystallography 21, 67–71

67. Kabsch, W. (1993) Automatic Processing of Rotation Diffraction Data from Crystals of Initially Unknown Symmetry and Cell Constants. Journal of Applied Crystallography 26, 795–800

68. Kabsch, W. (2010) Integration, scaling, space-group assignment and post-refinement. Acta Crystallogr D 66, 133–144

69. Kabsch, W. (2010) Xds. Acta Crystallogr D 66, 125–132

70. Bunkóczi, G., Echols, N., McCoy, A. J., Oeffner, R. D., Adams, P. D., and Read, R. J. (2013) Phaser.MRage: automated molecular replacement. Acta Crystallographica Section D-Structural Biology 69, 2276–2286

71. Evans, P. R. (2011) An introduction to data reduction: space-group determination, scaling and intensity statistics. Acta Crystallogr D 67, 282–292

72. Evans, P. R., and Murshudov, G. N. (2013) How good are my data and what is the resolution? Acta Crystallogr D 69, 1204–1214

73. Liebschner, D., Afonine, P. V., Baker, M. L., Bunkoczi, G., Chen, V. B., Croll, T. I., Hintze, B., Hung, L. W., Jain, S., McCoy, A. J., Moriarty, N. W., Oeffner, R. D., Poon, B. K., Prisant, M. G., Read, R. J., Richardson, J. S., Richardson, D. C., Sammito, M. D., Sobolev, O. V., Stockwell, D. H., Terwilliger, T. C., Urzhumtsev, A. G., Videau, L. L., Williams, C. J., and Adams, P. D. (2019) Macromolecular structure determination using X-rays, neutrons and electrons: recent developments in Phenix. Acta Crystallographica Section D-Structural Biology 75, 861–877

74. Read, R. J., and McCoy, A. J. (2011) Using SAD data in. Acta Crystallogr D 67, 338–344

75. Afonine, P. V., Grosse-Kunstleve, R. W., Echols, N., Headd, J. J., Moriarty, N. W., Mustyakimov, M., Terwilliger, T. C., Urzhumtsev, A., Zwart, P. H., and Adams, P. D. (2012) Towards automated crystallographic structure refinement with phenix.refine. Acta Crystallographica Section D-Structural Biology 68, 352–367

76. Emsley, P., and Cowtan, K. (2004): model-building tools for molecular graphics. Acta Crystallographica Section D-Structural Biology 60, 2126–2132

77. Terwilliger, T. C., Grosse-Kunstleve, R. W., Afonine, P. V., Moriarty, N. W., Zwart, P. H., Hung, L. W., Read, R. J., and Adams, P. D. (2008) Iterative model building, structure refinement and density modification with the wizard. Acta Crystallogr D 64, 61–69

78. Schindelin, J., Arganda-Carreras, I., Frise, E., Kaynig, V., Longair, M., Pietzsch, T., Preibisch, S., Rueden, C., Saalfeld, S., Schmid, B., Tinevez, J. Y., White, D. J., Hartenstein, V., Eliceiri, K., Tomancak, P., and Cardona, A. (2012) Fiji: an open-source platform for biological-image analysis. Nat Methods 9, 676–682

