## Supplemental Materials for "Atg23 Interacts With Both the N- and C-termini of Atg9 Via a Hydrophobic Binding Pocket"

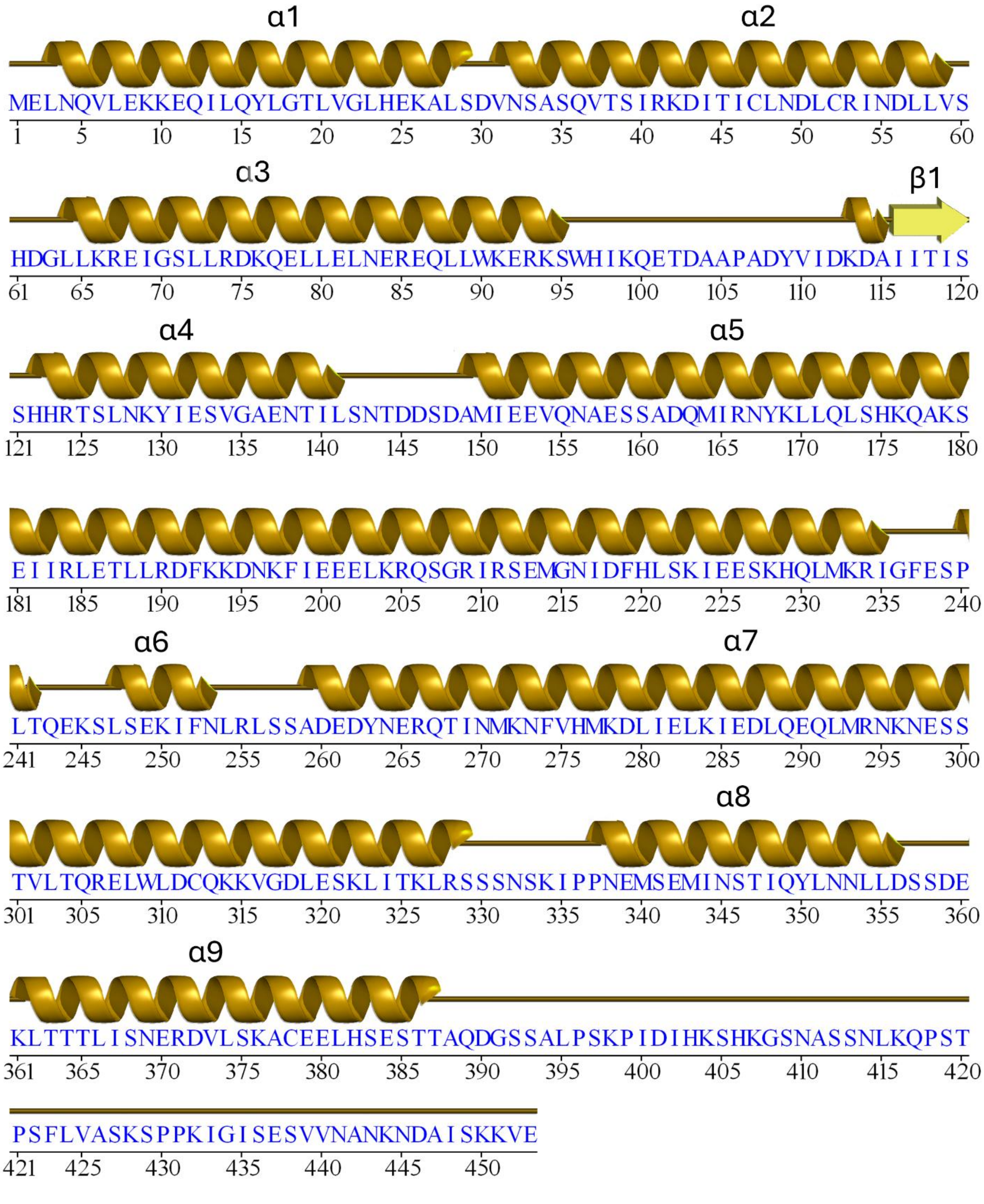


**Supplemental Figure 1. Secondary structure of the Atg23 AlphaFold 3 dimer model.** The amino acid sequence for Atg23 from Saccharomyces cerevisiae is shown with all elements of secondary structure from the AlphaFold 3 model indicated about the sequence and labeled. This figure was generated with PDBsum webserver (1).

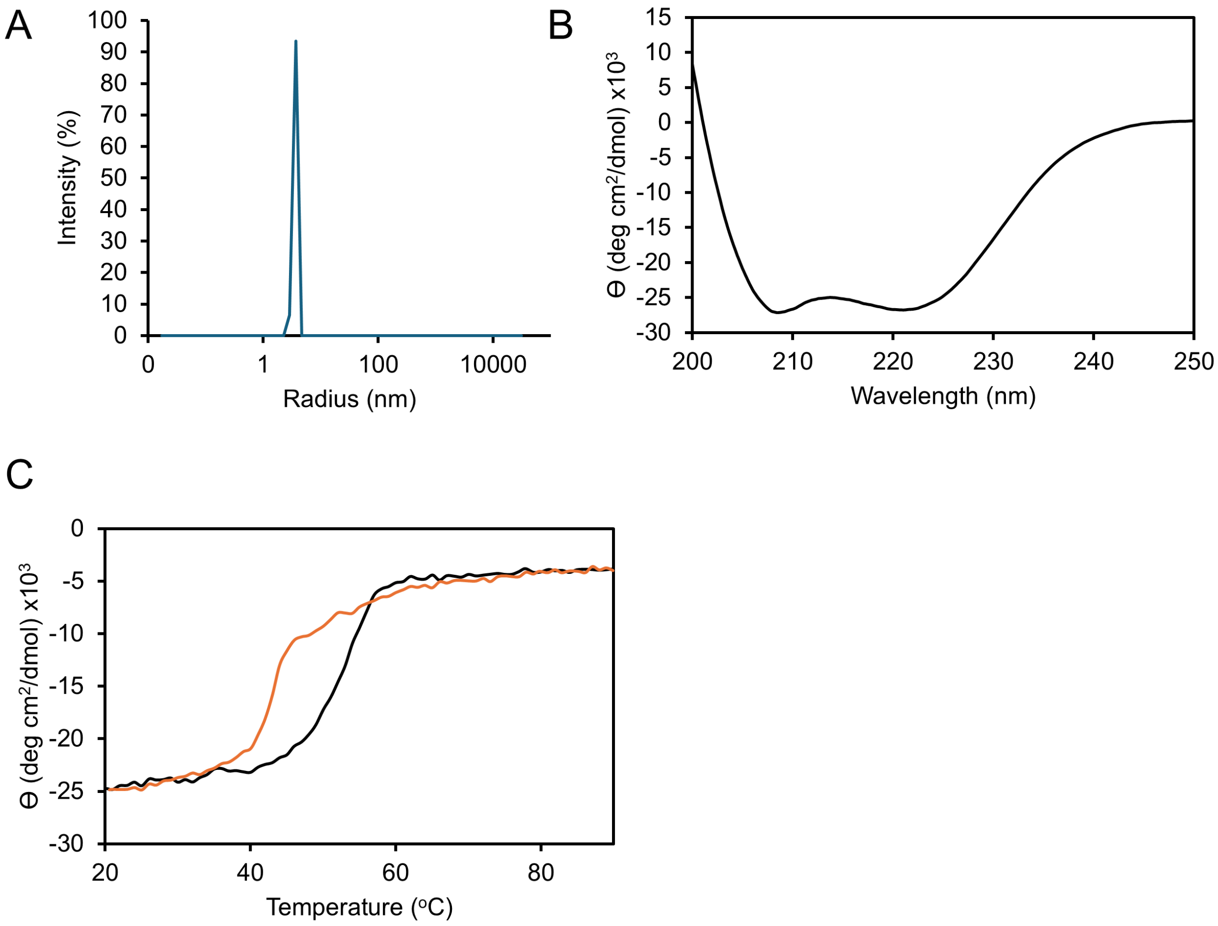


**Supplemental Figure 2. Additional validation of Atg23$_{ANNS}$. A)** Dynamic light scattering data of Atg23$_{ANNS}$. **B)** Circular dichroism data for Atg23$_{ANNS}$. **C)** Melting curves of Atg23 (orange) and Atg23$_{ANNS}$ (black) by monitoring the circular dichroism signal at 222 nm while the temperature of the cell was increased.

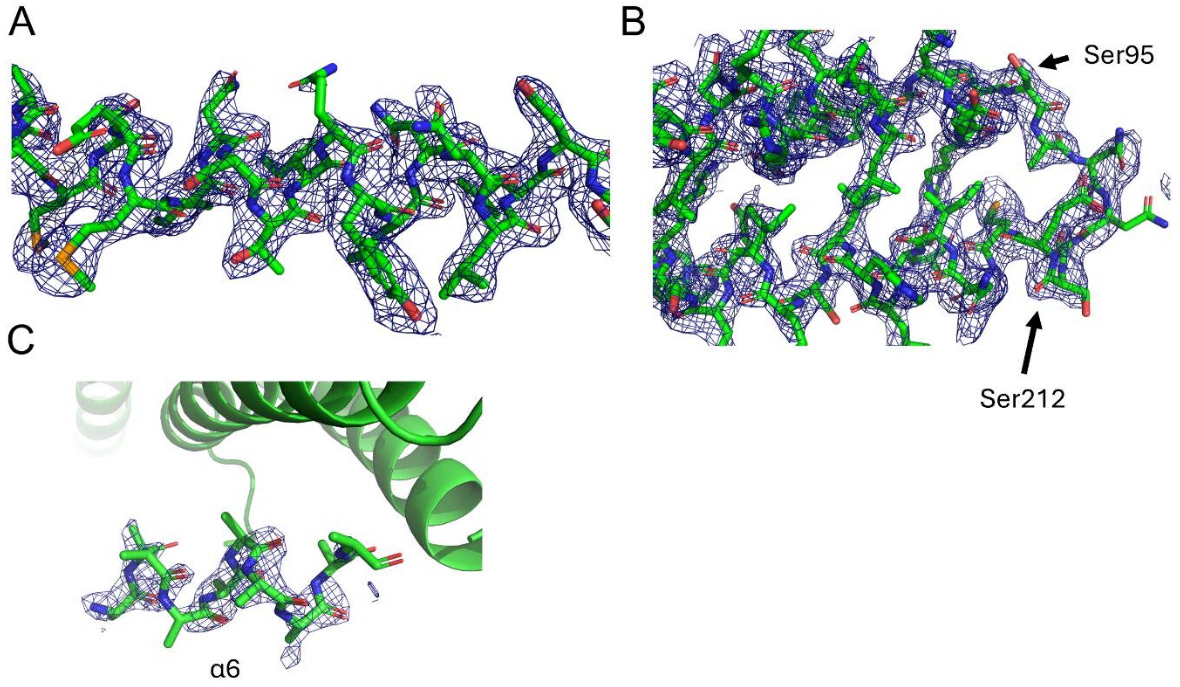


**Supplemental Figure 3. Representative Electron Density Maps. A)** 2Fo-Fc electron density map showing amino acids from α8 contoured at 1 ϴ. **B)** 2Fo-Fc electron density map showing the ANNS linker connecting Ser95 and Ser212 from Atg23 contoured at 1 ϴ. **C)** Fo-Fc omit electron density map contoured at 2.0 ϴ showing the weak alpha helical density that is likely α6.

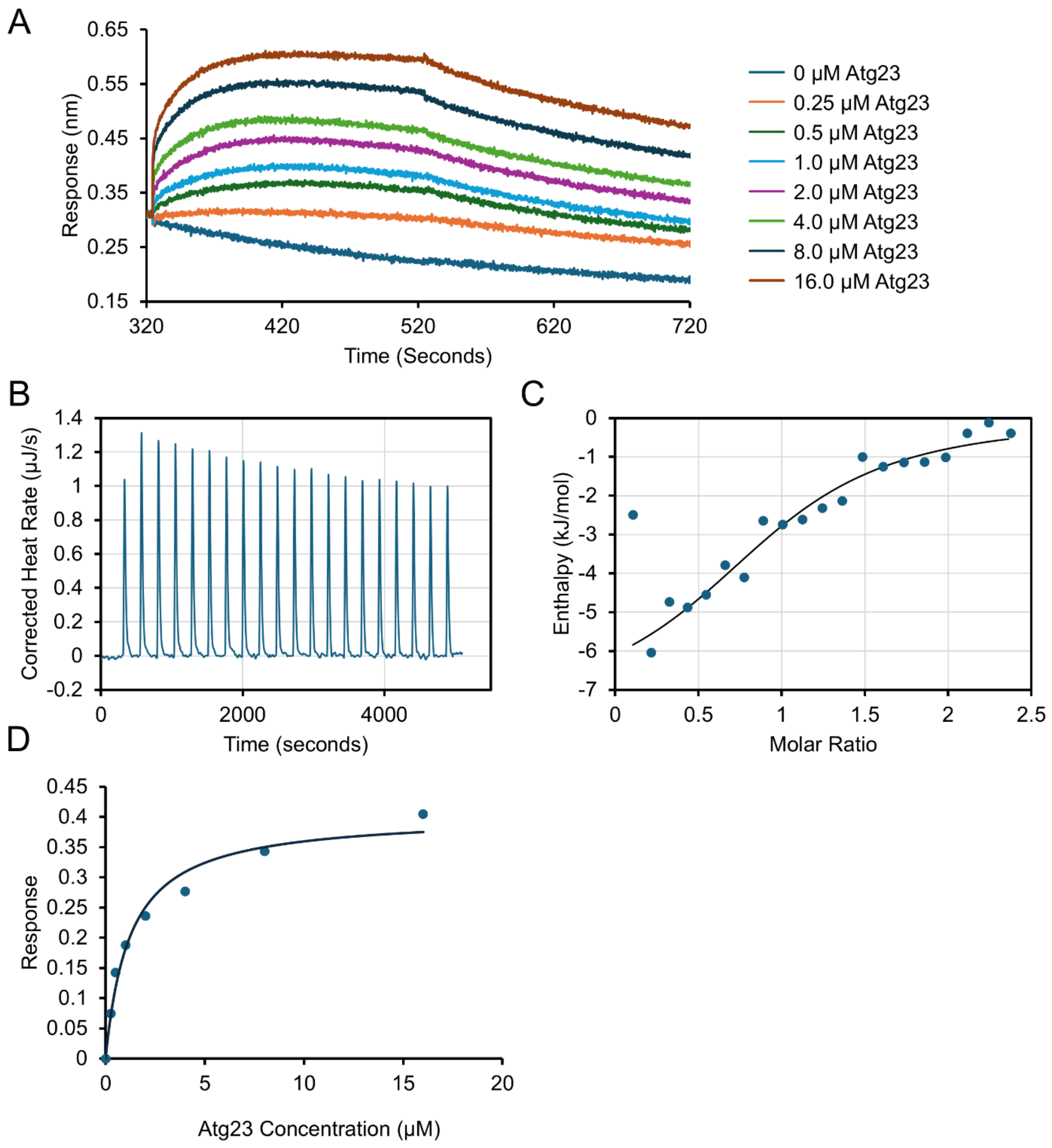


**Supplemental Figure 4. Additional binding data for the interaction between Atg23 and the N-terminus of Atg9. A)** Representative biolayer interferometry data after Atg9 1-255 was bound to the tip and then increasing concentrations of Atg23 were added from 0 to 16 μM. **B)** Representative raw thermogram from one of the triplicate isothermal titration calorimetry experiments where Atg9 1-255 was in the cell and Atg23 was titrated. **C)** Isotherm data from B where each integrated peak is shown as a blue dot and the fit to the data is shown as a black line. **D)** Steady state analysis of the BLI data for Atg9 1-48 and Atg23. Data is shown as dots and the fit to the data is shown as a line.

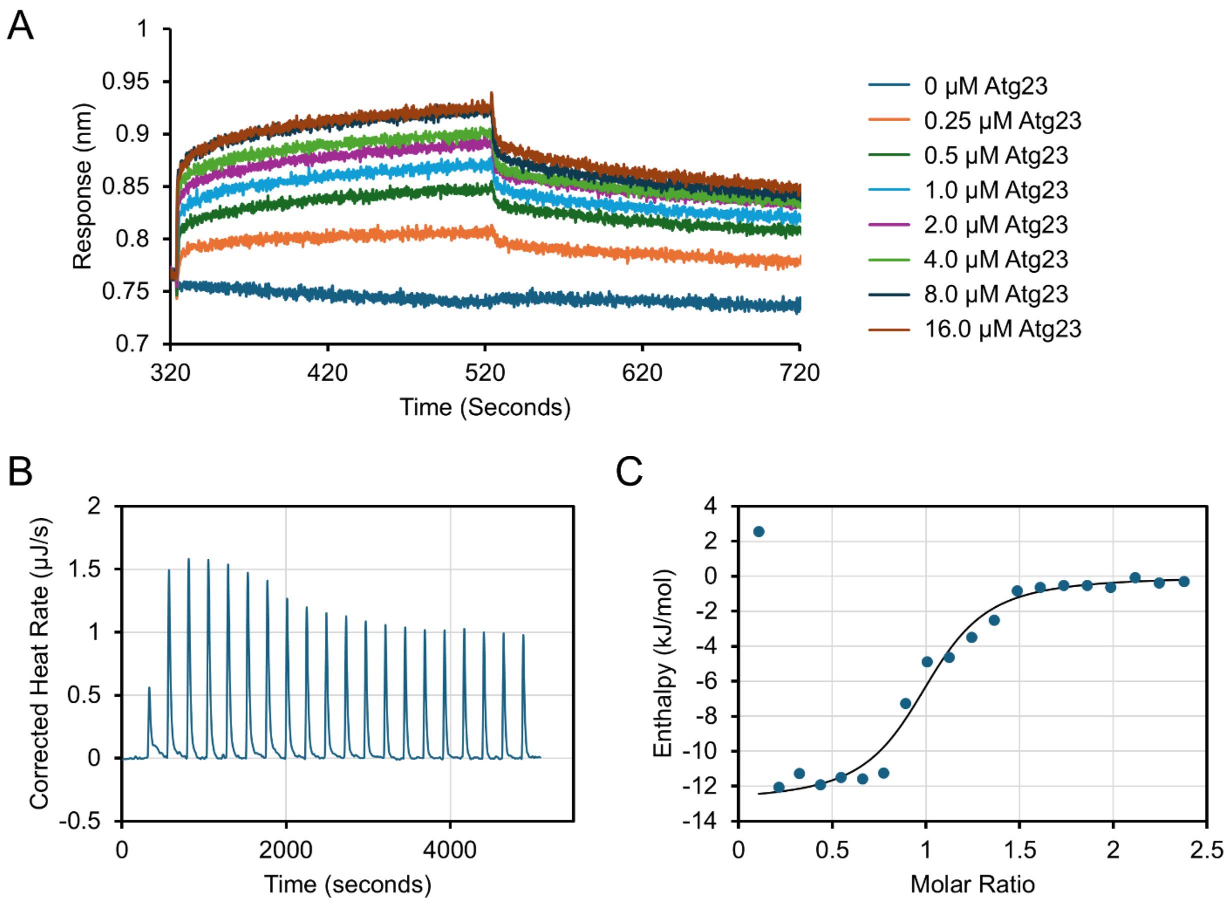


**Supplemental Figure 5. Representative binding data for the interaction between Atg23 and Atg9 864-997. A)** Representative biolayer interferometry data after Atg9 864-997 was bound to the tip and then increasing concentrations of Atg23 were added from 0 to 16 μM. **B)** Representative raw thermogram from one of the triplicate isothermal titration calorimetry experiments where Atg9 864-997 was in the cell and Atg23 was titrated. **C)** Isotherm data from B where each integrated peak is shown as a blue dot and the fit to the data is shown as a black line.

| Name | Vector | Details |
|---|---|---|
| Atg23 | pHis2 | 6XHis-TEV-Atg23 |
| Atg23 ANNS | pHis2 | 6XHis-TEV-Atg23$_{ANNS}$ |
| Atg23 L323R | pHis2 | 6XHis-TEV-Atg23 L323R |
| Atg23 M340E | pHis2 | 6XHis-TEV-Atg23 M340E |
| Atg23 A377E | pHis2 | 6XHis-TEV-Atg23 A377E |
| Atg23 A34E | pHis2 | 6XHis-TEV-Atg23 A34E |
| Atg23 Y16E | pHis2 | 6XHis-TEV-Atg23 Y16E |
| Atg9 1-255 | 1M | 6XHis-Atg9 1-255 -TEV-MBP |
| Atg9 1-48 | 1M | 6XHis-Atg9 1-48 -TEV-MBP |
| GST-Atg9 1-255 | pGST2 | 10XHis-GST-TEV-Atg9 1-255 |
| GST-Atg9 1-255 F17E | pGST2 | 10XHis-GST-TEV-Atg9 1-255, F17E |
| GST-Atg9 1-100 | pGST2 | 10XHis-GST-TEV-Atg9 1-100 |
| GST-Atg9 12-24 | 1G | 6XHis-GST-TEV-Atg9 12-24 |
| GST-Atg9 40-235 | pGST2 | 10XHis-GST-TEV-Atg9 40-235 |
| GST-Atg9 864-997 | pGST2 | 10XHis-GST-TEV-Atg9 864-997 |
| GST-Atg9 864-997 V982E | pGST2 | 10XHis-GST-TEV-Atg9 864-997, V982E |
| GST-Atg9 864-894 | pGST2 | 10XHis-GST-TEV-Atg9 864-894 |
| GST-Atg9 940-997 | pGST2 | 10XHis-GST-TEV-Atg9 940-997 |
| GST-Atg9 940-958 | pGST2 | 10XHis-GST-TEV-Atg9 940-958 |
| GST-Atg9 975-997 | pGST2 | 10XHis-GST-TEV-Atg9 975-997 |

Supplemental Table 1. A list of all the constructs used in this manuscript.

### Supplemental References

1. Laskowski, R. A., Jablonska, J., Pravda, L., Varekova, R. S., and Thornton, J. M. (2018) PDBsum: Structural summaries of PDB entries. *Protein Sci* **27**, 129-134
